# Loss of Complement Factor D suppresses alternative pathway activation but fails to reduce lipofuscin accumulation in the retinal pigmented epithelium of *Abca4^-/-^*mice

**DOI:** 10.1101/2025.10.25.684461

**Authors:** Bridget C. Ryan, Kaitlin E. Griffith, Caitlin N. Geczi, Katerina E. Kurz, Oliver Orchiston, Owen Smith, Xiaoya Xie, Adelaide H. Chow, Robert L. Chow

## Abstract

Stargardt disease (STGD1) is the most common inherited macular dystrophy, caused by loss-of-function mutations in *ABCA4* that result in bisretinoid-containing lipofuscin accumulation in the retinal pigment epithelium (RPE), and progressive photoreceptor degeneration. Oxidative stress and complement system activation have been implicated as contributors to disease pathogenesis, but the requirement for alternative pathway activation in STGD1 remains unclear. To directly assess this, we used a genetic approach to generate pigmented mice deficient for both *Abca4* and *Cfd*, an essential serine protease for alternative pathway initiation and amplification. Complement protein analysis revealed increased total C3 immunolabeling in the RPE and choroid of *Cfd^-/-^* mice, while C3d deposition at the RPE basal labyrinth and apical microvilli was markedly reduced, consistent with impaired alternative pathway activity. Western blotting confirmed altered C3 fragment profiles in *Cfd^-/-^* backgrounds, supporting a constitutive role for the alternative pathway in RPE complement activation. However, loss of *Cfd* did not prevent lipofuscin accumulation in the RPE of *Abca4^-/-^* mice. Under light-induced stress, we unexpectedly observed a modest attenuation of outer nuclear layer thinning in *Abca4^-/-^* that was unchanged by *Cfd* loss, which independently also showed a comparable rescuing effect. Together, these findings demonstrate that while the alternative pathway is a major driver of complement activation in the RPE and contributes only modestly to photoreceptor degeneration under light stress, its inhibition is insufficient to alter lipofuscin accumulation in pigmented *Abca4^-/-^*mice.

## Introduction

Stargardt disease (STGD1, MIM: 248200), also known as Stargardt macular dystrophy or juvenile macular degeneration, is the most common inherited macular dystrophy, with an incidence of approximately 1 in 8,000–10,000 (reviewed in references 1–4). It is an early-onset, progressive, autosomal recessive cone–rod dystrophy caused by loss-of-function mutations in *ABCA4*. Clinical manifestations typically include bilateral central vision loss, central scotoma, delayed dark adaptation, photosensitivity, photopsia, and abnormal color vision. Despite phenotypic variability, characteristic hallmarks often include yellow-white flecks, macular atrophy, and peripapillary sparing. A key diagnostic feature is the abnormal accumulation of lipofuscin in the retinal pigment epithelium (RPE), detectable by short wavelength (488 nm) autofluorescence. Disease progression involves RPE dysfunction and death, followed by secondary photoreceptor degeneration and disorganization. Current therapeutic strategies aimed at treating individuals with Stargardt disease include gene therapy, cell replacement, modulation of vitamin A transport, and targeting of the retinoid visual cycle; however, no approved treatments exist.

*ABCA4* encodes a membrane protein that localizes to photoreceptor outer segment discs and the retinal pigment epithelium (RPE) (5,6). It functions as an ATP-dependent flippase that transports the lipid–retinoid conjugate N-retinylidene-phosphatidylethanolamine (N-retinylidene-PE) from the luminal side of photoreceptor disc membranes to the cytoplasmic side (7). In the absence of functional *ABCA4*, N-retinylidene-PE accumulates and reacts with all-trans-retinal within the discs, leading to the formation of the bisretinoid fluorophore N-retinylidene-N-retinylethanolamine (A2E). A2E-containing outer segments are subsequently phagocytosed by the RPE, where bisretinoids accumulate as lipofuscin. Upon exposure to blue light, A2E undergoes photooxidation to form reactive epoxides (oxiranes) that induce oxidative stress and cell death in RPE cells *in vitro* (8,9). These processes are believed to underlie to chronic RPE dysfunction and photoreceptor degeneration characteristic of STGD1.

A2E photooxidation has been associated with increased activation of the complement system (10–12). Dysregulation of components regulating the alternative complement pathway has been reported in the RPE of both pigmented and albino *Abca4^-/-^* mice using immunostaining and Western blot analyses (13–16). The addition of rod outer segments from *Abca4^-/-^* mice to cultured human fetal RPE cells increased expression of complement regulatory proteins (CRPs) and oxidative stress markers, raising the possibility that A2E accumulation triggers complement activation through oxidative stress (16). Downregulation of CRPs in *Abca4^-/-^*mice may therefore enhance complement activation and oxidative damage in the RPE, contributing to cellular dysfunction and death. Furthermore, recombinant adeno-associated virus–mediated expression of a key mouse CRP, complement receptor 1-like protein y (CRRY), slowed photoreceptor degeneration and reduced both RPE autofluorescence at 488 nm and bisretinoid accumulation in *Abca4^-/-^*mice (15). Together, these findings highlight a critical link between A2E-induced oxidative stress, complement dysregulation, and RPE degeneration in STGD1.

Given the potential of complement inhibition as a therapeutic strategy for Stargardt disease, we sought to determine whether activation of the alternative pathway is required for disease progression in *Abca4^-/-^*mice. To block alternative pathway activation using a direct genetic approach, we generated *Abca4^-/-^*;*Cfd^-/-^* double mutants lacking complement factor D (CFD). CFD is a serine protease that cleaves factor B to form the C3 convertase and is therefore required to initiate the alternative pathway and for the amplification of all complement pathways. Complement activity, bisretinoid accumulation, and photoreceptor integrity were assessed using fundus autofluorescence imaging, histological and ultrastructural analyses, immunolabeling, and immunoblotting. This genetic loss-of-function approach represents an important proof of principle for testing the requirement of alternative pathway activation in RPE homeostasis and disease pathogenesis and informs the potential of complement-targeted interventions for inherited macular degeneration.

## Materials and Methods

### Mice

All experiments were conducted with ethics approval by the University of Victoria’s Animal Care Committee, following guidelines set by the Canadian Council for Animal Care. We generated experimental mice having four different genotypes: wild type, *Cfd^-/-^*, *Abca4^-/-^*, and *Abca4^-/-^;Cfd^-/-^* on a hybrid C57BL/6 X 129S1 background. Mice with the *Abca4* mutant allele (17) were derived from *Abca4^-/-^;Rdh8^-/-^* mice purchased from Jackson Labs (030503). *Cfd^-/-^* mice (18) were initially on a C57BL/6 background (gifted from Li Qiang, Columbia University). *Cfd^-/-^* mice were backcrossed into the 129S1 strain (Jackson labs, 002448). *Cfd^+/-^* and *Abca4^-/-^;Cfd^+/-^*mice were generated for experimental crosses, to produce both *Cfd^+/+^*and *Cfd^-/-^* littermate controls with and without *Abca4*. All experimental mice were confirmed by sequencing or genotyping to be homozygous for the Leu450 allele of *Rpe65*, which is associated with increased sensitivity to photoreceptor degeneration (19–21). Sequencing was also performed to confirm that mice did not carry the *Rd8* mutation in *Crb1,* which sometimes appears in vendor mouse lines and can confound ocular phenotypes (22). *C3* KO mice (Jackson Labs, 029661) were included as a negative control in C3 immuno-assays.

### Genotyping and Sequencing

Genotyping and sequencing were performed using genomic DNA (gDNA) prepared from mouse ear biopsies digested in 0.05 M NaOH for 10 minutes at 98°C, followed by neutralization with 0.5 M Tris pH 8.0 to a final concentration of 0.125 M. All genotyping PCRs were performed using KAPA2G Fast Hotstart genotyping mix (Millipore Sigma, kk5621). Primers are listed in Table 1. PCR amplification for sequencing of *Cfd* and *Rpe65* were performed Taq DNA polymerase (Froggabio, T-5000) and Phusion High-Fidelity DNA Polymerase (NEB, M0530S), respectively. PCR products used for sequencing were purified using QIAquick PCR purification kit (Qiagen, 28104) and sequenced (Eurofins Genomics, Canada).

**Table 1.**
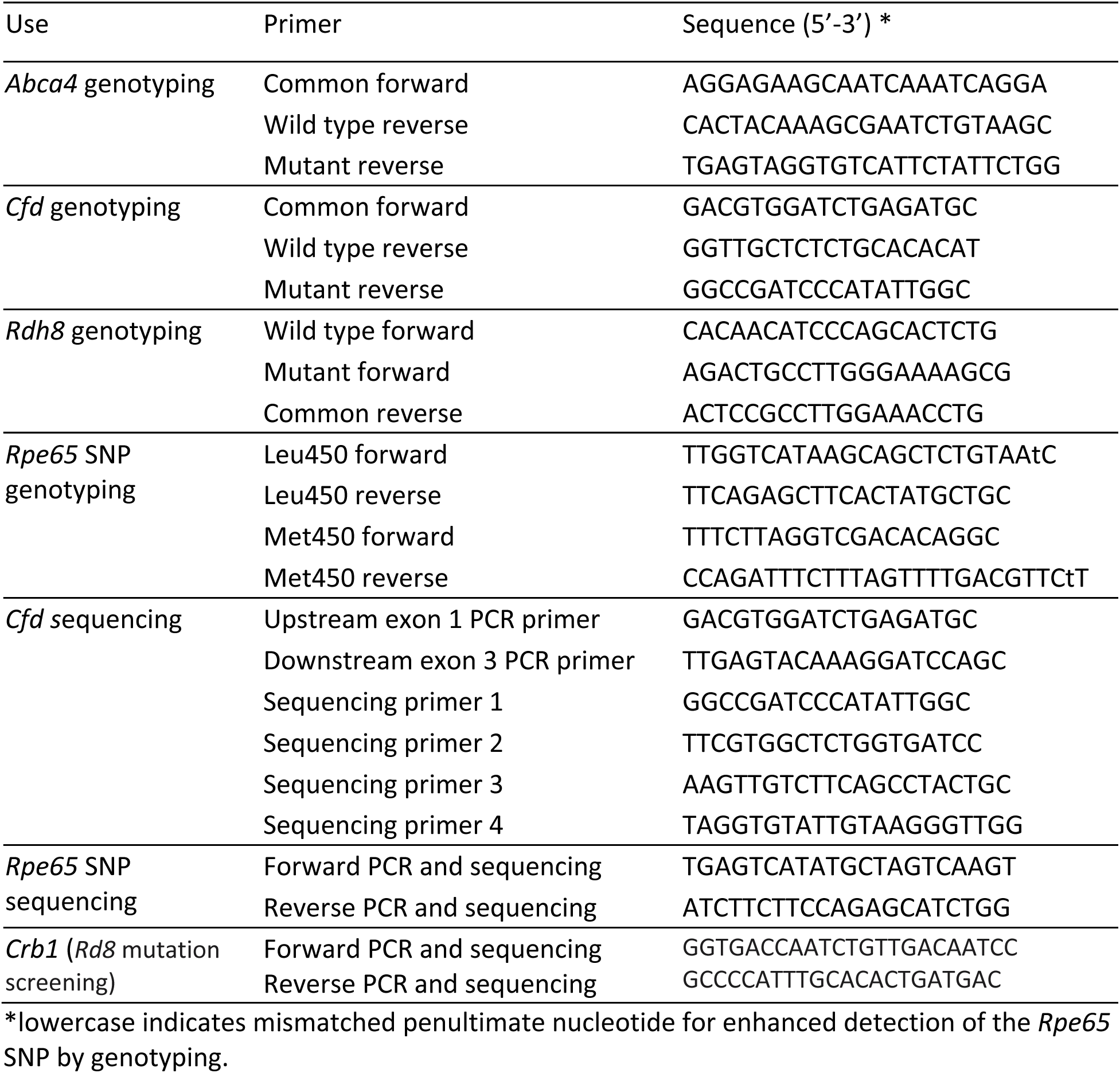
Summary of genotyping and sequencing primers.

*Abca4* and *Rdh8* genotyping: Thermocycling conditions used the hot start method with 94°C melt for 30 seconds, 65°C annealing for 30 seconds, 72°C extension for 30 seconds for the first cycle, with the annealing temperature dropping by 1°C each cycle for five cycles, finishing with an annealing temperature of 60°C for 30 cycles. *Cfd* genotyping: Thermocycling conditions used 94°C melt for 45 seconds, 58°C annealing for 45 seconds, 72°C extension for 45 seconds, repeating for 32 cycles.

*Rpe65* SNP genotyping: Thermocycling conditions used the hot start method with 94°C melt for 30 seconds, 58°C annealing for 30 seconds, 72°C extension for 30 seconds for the first cycle, with the annealing temperature dropping by 1°C each cycle for five cycles, finishing with an annealing temperature of 53°C for 32 cycles. *Cfd* mutation sequencing: Thermocycling conditions used the hot start method with 94°C melt for 30 seconds, 62°C annealing for 30 seconds, 72°C extension for 90 seconds for the first cycle, with the annealing temperature dropping by 1°C each cycle for five cycles, finishing with an annealing temperature of 57°C for 30 cycles. *Rpe65* codon 450 SNP sequencing: Thermocycling conditions used 98°C melt for 15 seconds, 61°C annealing for 30 seconds, 72°C extension for 15 seconds, repeating for 35 cycles.

### Eye tissue harvest and fixation

Mice were dark adapted for 20-24 hours prior to tissue harvest or exposed to light stress as described below. Tissue dissections were performed at the same time each morning. Tissue harvested for 488 nm lipofuscin quantification was performed in the dark under dim red light. Mice were anesthetized by inhalant isoflurane (AVP, 8061652) and euthanized by cervical dislocation. Enucleated eyes were processed for either Western blot, flat mount or tissue sectioning. For Western blot and flat mount preparation, the anterior chamber, lens, retina, and extraocular tissue were removed. RPE, choroid and sclera were either flash frozen on liquid nitrogen for Western blot or butterflied and fixed flat in 1X phosphate buffered saline (PBS) with 4% paraformaldehyde (PFA, Electron Microscopy Sciences, 157-8) for one hour at room temperature for flat mount. Eyes used for preparing tissue sections were punctured with a 27 G needle at the nasal limbus and fixed in 1X PBS with 4% PFA for one hour at room temperature. Following fixation, whole eyes were marked with a suture at the nasal side using a microsurgical needle with suture thread (FST, 12051-08).

### Tissue section preparation

Eyes were sectioned through the optic nerve head (ONH) along the inferior-superior axis.

For paraffin-embedded sections, sutured eyes were dehydrated in 70% ethanol. Paraffin embedding and preparation of 4 µm thick sections was performed by Wax-it Histology Services (Vancouver, Canada). For frozen sections, eyes were cryoprotected in 1X PBS with 20% sucrose, embedded in OCT medium (EMS, 62550) and sectioned at a thickness of 12 µm.

### Immunolabeling

Paraffin sections were deparaffinized though successive washes twice for 5 minutes in 100% xylenes, twice for 3 minutes in 100% ethanol, once for 1 minute in 95% ethanol and once for 1 minute in 80% ethanol. Antigen retrieval was performed on deparaffinized sections using the steamer method previously described (23) and a 5.5-quart digital steamer (Hamilton Beach, 37530C), by incubating slides in sodium citrate buffer (10 mM Tri-sodium citrate, 0.5% tween- 20, pH 6.0) at 100°C for 30 minutes. Frozen and paraffin sections were permeabilized in 1X PBS with 1% Triton X-100 for 10 minutes at room temperature. Flat mount eyecup tissue was permeabilized for one hour at room temperature in PBS with 1% Triton X-100 prior to immunolabelling. Tissue sections were incubated overnight at 4°C with primary antibodies diluted in 1X PBS (Table 2). Anti-C3 immunofluorescence was performed using frozen sections and anti-C3d immunofluorescence using paraffin sections. Detection was performed at room temperature for 90 minutes using Alexa647-conjugated secondary antibodies (Table 2) to minimize the contribution of RPE autofluorescence to the immunofluorescence signal. Slides were mounted using Immu-Mount (Shandon, 9990402).

**Table 2.** Summary of antibodies used for fluorescence microscopy.

| Antibody | Dilution | Supplier | Product # | Use |
| --- | --- | --- | --- | --- |
| rabbit anti- ZO-1 | 1:500 | ThermoFisher | 67-7300 | AF RPE marker |
| goat anti-mouse C3 | 1:100 | MP Biomedicals | 55463 | IF frozen |
| goat anti-mouse C3d | 1:100 | R&D Systems | AF2655 | IF paraffin |
| rabbit anti-RPE65 | 1:400 | ThermoFisher | PA5-110315 | IF paraffin |
| donkey anti-rabbit Alexa 647 | 1:500 | Invitrogen | D1306 | IF secondary |
| donkey anti-goat Alexa647 | 1:500 | Invitrogen | A21447 | IF secondary |

### 488 nm autofluorescence quantification

488 nm autofluorescence was quantified from 6-mo and 12-mo paraffin eye sections and from 12-mo flat mount eyecup tissue. Imaging was performed using 488 nm excitation and 525 nm ± 25 nm emission on a Nikon C2+ confocal microscope. Tissue sections and flat mount eyecups were imaged using 60X oil and 20X objectives, respectively. Flat mount images were collected as multiple z-stacks, spaced 1 µm apart. Samples analyzed as part of the same experiment were imaged using the same imaging settings, which were established to minimize oversaturated pixels in positive control samples. For tissue sections, eight non-overlapping images were collected per sample, with four images collected across two tissue sections, two images directly superior and two directly inferior to the ONH. For flat mount tissue, four images were collected per sample, one from each quadrant of the central eye cup. A uniform fluorescent slide (Chroma, 92001) was also imaged for performing flat field correction. Mean fluorescence intensity was quantified from the RPE using FIJI ImageJ software. Prior to performing quantification of fluorescence intensity, images were blinded and randomized, flat field corrected, and maximum intensity projected using a custom ImageJ macro. For flat mount tissue, 488 nm autofluorescence was only quantified from regions with in-focus ZO-1 labelling. The ZO-1 labelling was also used to define individual RPE cells as regions of interest using a threshold mask in ImageJ, for quantification of fluorescence intensity from individual RPE cells. For each sample, the mean fluorescence intensity in the RPE was averaged across replicate images to avoid pseudo-replication.

### C3 immunofluorescence quantification

C3 protein levels in RPE and choroid were quantified from eye tissue sections by indirect immunofluorescence. For each sample, a tissue section lacking primary antibody was included as a negative control, to assess the contribution of general tissue fluorescence. Eye sections through the optic nerve head were used, and sample genotypes were anonymized prior to staining and confocal imaging using a 60X oil objective. Multiple non-overlapping replicate images were collected per sample, two images immediately superior and two inferior to the optic nerve head. Imaging settings were established using samples having the highest immunofluorescence signal to maintain pixels below saturation. All images used for a single experiment were imaged using the same settings. General tissue autofluorescence at 488 nm was also included for identifying tissues of interest for imaging and quantification. Prior to image processing and analysis, all images associated with a single experiment were blinded and randomized using a custom ImageJ macro. RPE and choroid were defined as regions of interest using 488 nm autofluorescence and mean immunofluorescence pixel intensity was measured using ImageJ. Data were analyzed by averaging the mean immunofluorescence intensity across replicate images from the same sample and subtracting the mean intensity measured from the no-primary control.

### Western blot

Western blots were performed using eyecup tissue (RPE, choroid, sclera) from one eye per mouse. Eyecup lysate was prepared in T-PER Tissue Protein Extraction Reagent (Thermo Fisher, 78510) with protease and phosphatase inhibitor cocktail (1:100, Thermo Fisher, 78440) and 0.01 M EDTA. Tissues were homogenized using a Disruptor Genie (Scientific Industries, SI- DD38) at 1000 RPM for 1 minute at 4°C. Following homogenization, lysate was incubated at 4°C for 1 hour on a rocker. Cell debris was removed by centrifugation at 10,000 g for 5 minutes at 4°C. One-quarter of the total eyecup lysate was used for a single gel lane and excess unreduced lysate was stored at -80°C. Lysate was reduced in 1X lithium dodecyl sulfate (LDS) buffer (Thermo Fisher, NP0007) and 1X sample reducing agent (Thermo Fisher, NP0004) for 12 minutes at 75°C and was immediately loaded into wells of NuPAGE 4-12% Bis-Tris gels (Thermo Fisher, NP0321BOX) in NuPAGE 3-(N-morpholino) propanesulfonic acid (MOPs) SDS running buffer (Thermo Fisher, NP0001). Gels were run on ice at 150 V for 120 minutes. The MagicMark XP Western Protein Standard (Thermo Fisher, LC5602) was used as a protein ladder. Proteins were wet transferred onto a polyvinylidene fluoride (PVDF) membrane (Millipore Sigma, IPFL07810) in 1X NuPAGE Transfer buffer (Thermo Fisher, NP00061) with 10% ethanol at 20 V for 100 minutes. Membranes were blocked in Intercept PBS Blocking Buffer (LI-COR Biosciences, 927-70001). Antibodies were diluted in blocking buffer with 0.15% Tween-20. Membranes were incubated at 4°C for 15 hours in primary antibody solution and 1 hour in secondary solution at room temperature. Membranes were washed three times in 1X PBS with 0.1% tween-20 after each antibody incubation. The antibodies used for Western blotting and their dilutions are listed in Table 3. When quantification of both RPE65 and β-actin were performed from the same blot, labelling and imaging for RPE65 was performed first, followed by β-actin. Membranes were scanned using an Odyssey ClX scanner (LI-COR Biosciences, model no. 1214) and Image Studio Lite acquisition software version 5.2.

**Table 3.** Summary of antibodies used for Western blot.

| Antibody | Dilution | Supplier | Product # | Use |
| --- | --- | --- | --- | --- |
| goat anti-mouse C3d | 1:2000 | R&D Systems | AF2655 | WB primary |
| mouse anti-RPE65 | 1:1500 | Novus | NB100-35555 | Loading control |
| mouse anti-β-actin | 1:1250 | Genscript | A00702 | Loading control |
| donkey anti-goat 800 CW | 1:15,000 | LI-COR | 926-32214 | WB secondary |
| donkey anti-mouse 680 RD | 1:20,000 | LI-COR | 926-68072 | WB secondary |

Quantification of relative C3 fragment protein levels was performed using blots labelled with the anti-C3d antibody and Empiria Studio software version 2.3.0.154. Relative levels of C3 preprotein, C3 α-chain (C3α), opsonized C3b α-chain, iC3(H2O) α72, iC3b α62 and the C3dg α- chain were first normalized to combined relative levels of both RPE65 and β-actin, calculated by geometric mean. To control for inter-blot signal variability, lane normalized C3 fragment quantities were then normalized to wild type C3α within each blot.

### Light-Induced Photoreceptor Degeneration

We used a bright light model for inducing photoreceptor degeneration following a general approach described previously (19,24–31). Between 8:00 and 9:00 am, 24 hours before beginning light exposure, mice were habituated to holding containers for 15 minutes. Mice were then dark adapted by housing them in red plastic light-blocking cages (Tecniplast, GM500 Red X-Temp) placed behind an opaque black plastic curtain in a dark room (illumination = 0 lux), with access to food and water. Mice were monitored using a dim red-light headlamp (Petzl) (illumination < 2 lux). 24 hours after starting dark adaptation, Mydriacyl ophthalmic eyedrop solution (1% tropicamide, Alcon, ordered through AVP, 2102452) was applied to each eye of awake mice under dim red light. Mice were maintained in the dark in their home cages for 15 minutes prior to light exposure. Light exposure was performed by housing mice individually in holding containers (Supplementary Figure 1) placed under a fluorescent light apparatus (Fusion bright 2 ft X 6 144 W T4 fluorescent lights, colour temperature 6500 K from Grow Lights Canada, FLP26). Up to four mice were processed at one time. Prior to beginning each experiment, illumination inside the holding containers was confirmed to be 19-20 klux using a light meter (Extech instruments, LT300), with detector positioned facing upward inside the holding container. Fans were used to maintain temperatures between 22-23°C during the experiment. Experimental mice were exposed to light for 4 hours. Following light exposure, mice were dark adapted for 3 or 6 days prior to tissue harvest.

### Outer Nuclear Layer Thickness Quantification

Outer nuclear layer (ONL) thickness was measured across the entire inferior to superior retina from all experimental genotypes using DAPI-labelled frozen tissue sections with and without light stress. Images were collected using the 20X objective and five z-stacks spaced 2 µm apart were collected per image. Two replicate tissue sections were imaged per mouse.

Image analysis was performed using FIJI ImageJ. Maximum intensity projected images collected from the same tissue section were stitched into composite images using the pairwise stitching plug-in in ImageJ (32). Composite images were blinded and randomized using a custom ImageJ macro prior to analysis. To measure ONL thickness, a segmented line was first drawn through the middle of the ONL, spanning the proximal to distal retina on both the inferior and superior side of the optic nerve head, for each blinded composite image. Segmented lines were marked at 250 µm intervals, starting at the optic nerve head, using a custom ImageJ macro. The average ONL thickness within each 250 µm interval was calculated by measuring the ONL area between each interval in um^2^ and dividing it by the interval length. ONL thickness measurements within each interval were averaged between replicate tissue sections from the same mouse to avoid pseudo-replication.

### Statistics

GraphPad Prism was used to generate graphs and perform all statistical analyses.

Statistical test, p values, and the number of biological replicates are indicated in the results and figure legends for each graph.

## Results

To examine the role of CFD in the progression of the *Abca4^-/-^*mutant phenotype, we generated *Abca4^-/-^*; *Cfd^-/-^*mice. Mice were genotyped to ensure that they carried the Leu450 allele of Rpe65 which has been shown to have higher isomerase activity, lipofuscin levels and increased sensitivity to light induced retinal damage than the Met450 variant (19–21). We first examined RPE 488 nm autofluorescence as a proxy for bisretinoid accumulation. Flat-mount eye cup preparations from 12-month-old mice were used to maximize the area of RPE that could be quantified (10–12). A >2-fold increase in RPE autofluorescence in *Abca4^-/-^* mice relative to wild type was observed when mean 488 nm autofluorescence was measured across all imaged regions of eyecup (Figure 1B) and within individual RPE cells (Figure 1C). This increase in autofluorescence was similarly observed in *Abca4^-/-^*; *Cfd^-/-^* mice and indistinguishable from *Abca4^-/-^*mice. In *Cfd^-/-^* mice, RPE 488 nm autofluorescence was comparable to that observed in wild type mice. Mice were also examined at 6-months of age, but elevated RPE 488 nm autofluorescence was not observed in *Abca4*^-/-^ mice relative to wild type (Supplementary Figure 2). Together, these findings suggest that CFD-mediated initiation of the alternative pathway and progression of the C3 convertase amplification loop do not play a major role in regulating RPE bisretinoid levels in pigmented mice.

**Figure 1.**
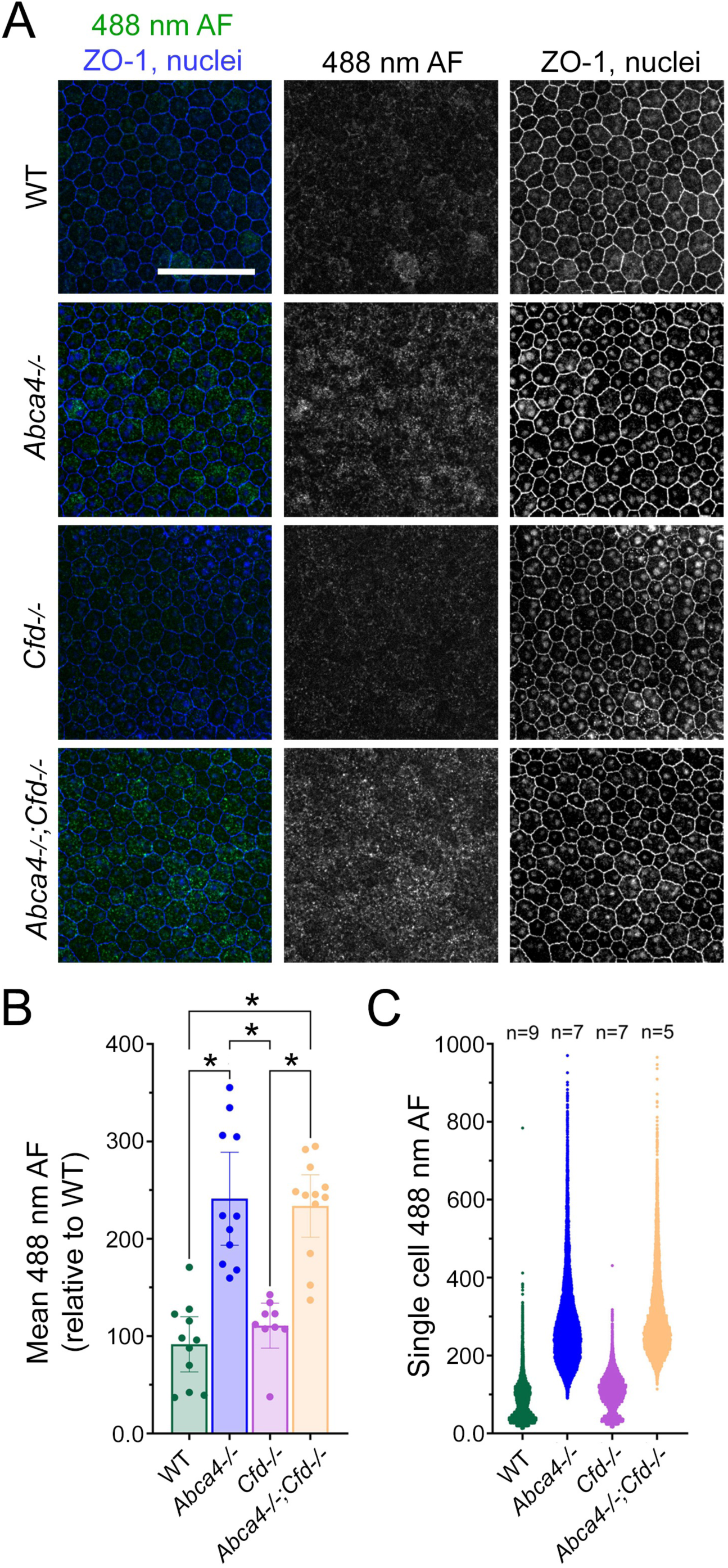
Loss of *Cfd* does not rescue the lipofuscin accumulation phenotype in 12-mo *Abca4^-/-^* mutant RPE. A) Representative images showing 488 nm autofluorescence (AF, green) in 12-mo wild type, *Abca4^-/-^*, *Cfd^-/-^* and *Abca4^-/-^*;*Cfd^-/-^*dark-adapted and PFA-fixed flat mount RPE. ZO-1 immunofluorescence and nuclear labelling are shown in blue. B) Quantification of mean 488 nm AF from flat mount RPE IMAGES. Each data point represents the average of four separate replicate images collected from one mouse and data are represented at mean +/- 95.% C.I. Replicate images were collected from the central eyecup, one image per eyecup quadrant. Data were analyzed by one-way ANOVA with Tukey’s multiple comparisons test. * indicates p<0.0001. C) Quantification of 488 nm AF in single RPE cells. Data are presented with each point representing the mean 488 nm AF in a single RPE cell. Approximately 1,500-3,000 cells were quantified per individual mouse.

### As changes in C3 fragment quantities have previously been reported in the RPE of

*Abca4^-/-^* mice (13,14,16,33), we next examined C3 immunofluorescence labeling in the RPE of *Abca4^-/-^* and *Abca4^-/-^; Cfd^-/-^* mice. C3 labeling was quantified in mouse eyecup structures, such as the RPE and choroid, using antibodies generated against full-length mouse C3 protein (referred to as “anti-C3”, MP Biomedicals) or purified mouse C3d (referred to as “anti-C3d”, R&D Systems) (Figure 2). In wild type sections, anti-C3 labelling (Figure 2A) was detected primarily along the basal side of the RPE, with most of the labelling suspected to be adjacent to the basal RPE in Bruch’s membrane and the choroid. Anti-C3d labelling (Figure 2B) was detected robustly at the basal side of the RPE, and faint labelling was also detected at the apical side of the RPE and bands within the RPE. Phalloidin labeling of F-actin and RPE65 labeling of the RPE somata (34) revealed that anti-C3d immunolabeling is primarily associated with the RPE basal labyrinth and weakly associated with the RPE apical microvilli (Supplementary Figure 3A,B). Phalloidin staining also revealed that 488 nm autofluorescence can be used reliably to localize these RPE structures, and to define the RPE for subsequent quantitative immunolabeling experiments.

**Figure 2.**
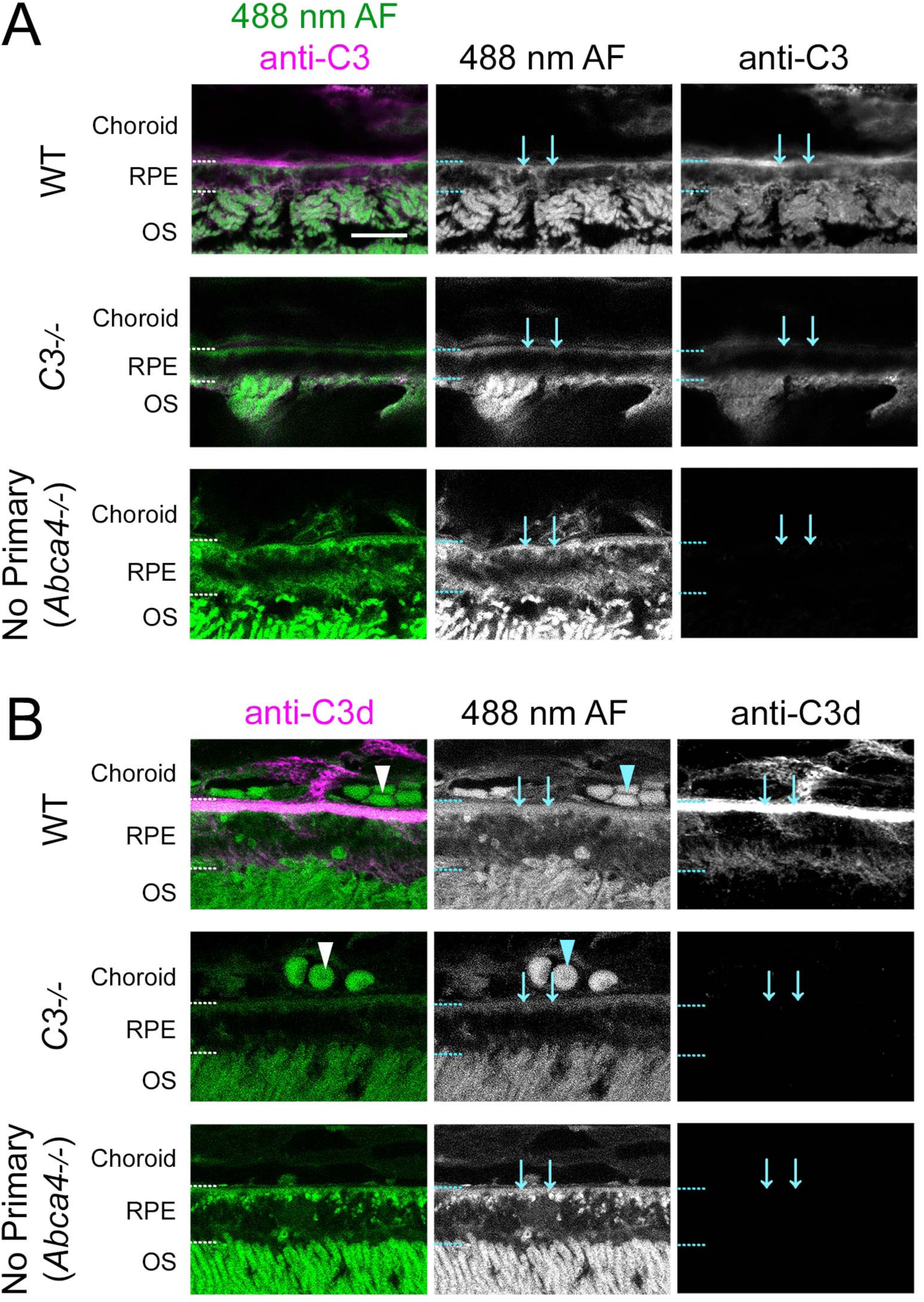
Specificity of immunofluorescence labeling produced by different C3 antibodies in the mouse RPE of wild type and negative control samples. Immunofluorescence labeling of PFA-fixed wild type and *C3^-/-^* mouse eyecup tissue with A) anti-C3 (MP Biomedicals, 55463) and B) anti-C3d (R&D Systems, AF2655) antibodies, showing genuine antibody labeling in the RPE of wild type samples, localized primarily to the basal side of the RPE. Note suspected non-specific binding of the anti-C3 primary antibody to photoreceptor outer segments (OS). *Abca4^-/-^*samples were included as a no primary control, to show general tissue AF. Labeling using the anti-C3 antibody was performed using frozen sections and labeling using the anti-C3d antibody was performed using paraffin-embedded tissue sections. General tissue AF at 488 nm was included to assist in identifying eyecup structures and for comparison to immunofluorescence signal in the no primary control conditions. Note that each C3 antibody may detect multiple C3 fragments. The scale bar represents 10 µm and is consistent across all images. Choroid, RPE and OS are shown. Arrowheads show red blood cells, which are fluorescent at 488 nm in paraffin sections. Arrows show the location of suspected RPE basal labyrinth.

Notably, the RPE basal labyrinth membrane can be seen as a band of autofluorescence at the basal side of the RPE, serving to differentiate RPE from adjacent Bruch’s membrane and choroid. Anti-C3d labelling was also observed associated with choroidal capillaries, as evidenced by the presence of autofluorescent red blood cells. ZO-1 labeling of RPE tight junctions revealed that C3d immunofluorescence is also weakly associated with lateral RPE cell membranes (Supplementary Figure 3C). The signal produced by both antibodies was largely absent from the RPE and choroid in the *C3^-/-^* and no primary negative controls (except for non-specific labeling of damaged photoreceptor outer segments with anti-C3), suggesting that the immunolabeling signal is specific.

We next performed quantitative immunofluorescence using the anti-C3 and anti-C3d antibodies to measure changes in the quantity of different C3 fragments in the RPE of *Abca4* and *Cfd* mutant mice (Figure 3). Anti-C3 signal in the RPE appeared generally elevated when *Cfd* was knocked out, particularly in *Abca4^-/-^* mice (Figure 3A, B). Additionally, C3 labeling appeared elevated in the choroid of *Cfd^-/-^* mice relative to *Cfd^+/+^*(Figure 3A). The effect of the *Cfd* mutation on total C3 immunofluorescence was especially pronounced when samples were grouped by *Cfd* genotype, with *Cfd^-/-^*mice showing elevated anti-C3 labeling relative to *Cfd^+/+^* (Supplementary Figure 4B). Anti-C3d immunofluorescence showed a pronounced loss of signal in the basal labyrinth, apical microvilli and lateral RPE when *Cfd* was mutated (Figure 3C, Supplementary Figure 4A). While in theory, the anti-C3d antibodies may be able to detect multiple C3 fragments (including intact C3) in PFA-fixed tissue sections, the loss of C3d labeling suggests that the C3d antibody recognizes cleaved C3 in immunolabeled fixed tissue.

**Figure 3.**
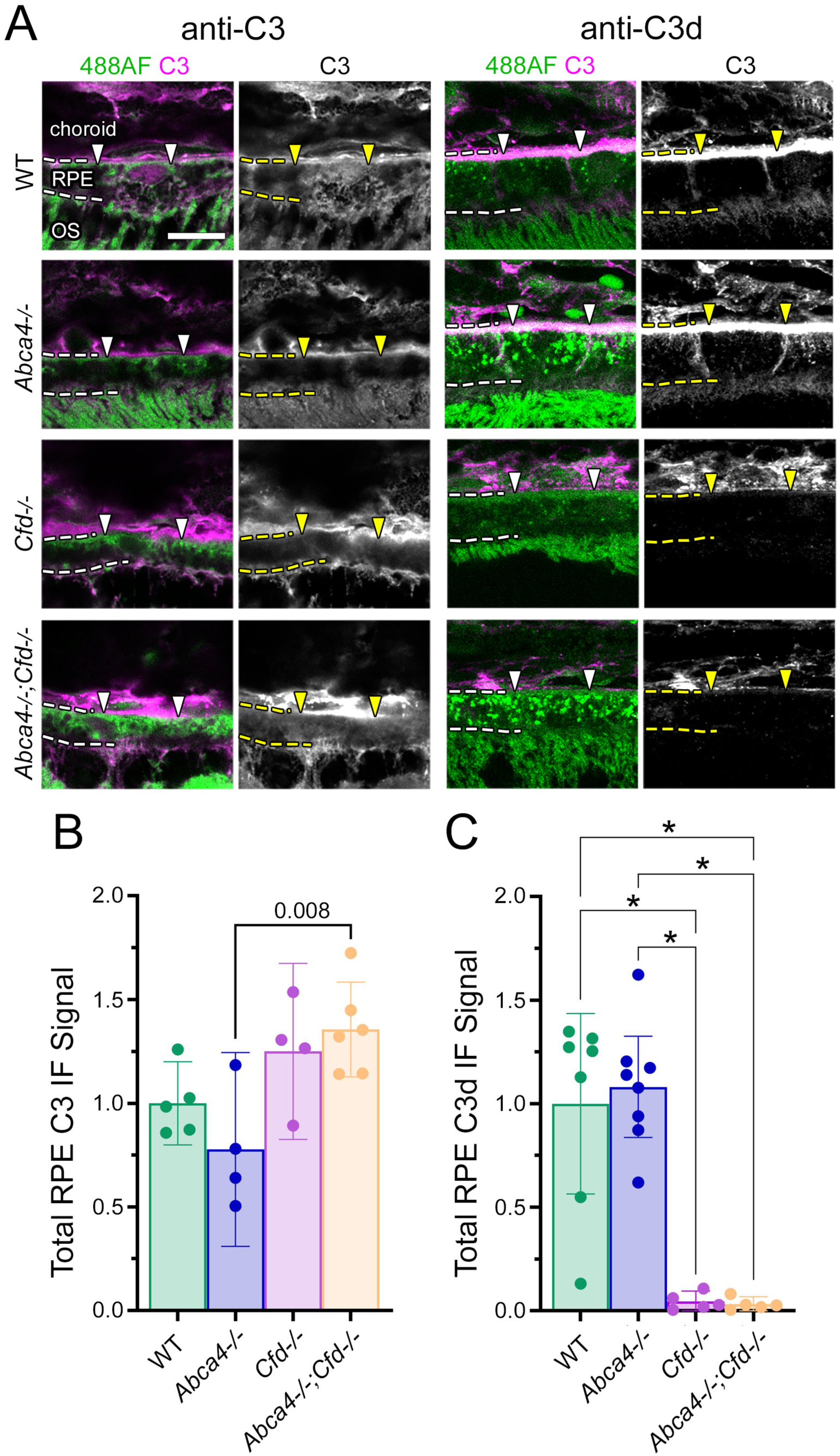
C3 fragment quantification by immunofluorescence in the RPE of *Abca4^-/-^* and *Cfd^-/-^* mice. A) Representative images showing immunofluorescence labelling in PFA-fixed wild type, *Abca4^-/-^*, *Cfd^-/-^*and *Abca4^-/-^*;*Cfd^-/-^* mouse eyecup tissue sections, using anti-C3 (MP Biomedicals) and anti-C3d (R&D Systems) antibodies. Anti-C3 immunofluorescence was performed using 6-12-mo frozen sections and anti-C3d immunofluorescence was performed using 12-mo paraffin embedded sections. For each image, 488 nm AF (green) and C3 immunofluorescence (magenta) are shown together, to aid in localizing the immunofluorescence signal. C3 immunofluorescence is also shown on its own in greyscale. Choroid, RPE and photoreceptor outer segments (OS) are labelled, with dotted lines indicating the basal (top) and apical (bottom) boundaries of the RPE. Arrowheads show the location of Bruch’s membrane, between the RPE basal labyrinth and the choroid region. Scale bar represents 10 µm. B-C) Quantification of total RPE immunofluorescence signal in each genotype using the anti-C3 and anti-C3d antibodies, respectively. Total immunofluorescence was quantified by measuring the mean immunofluorescence signal in the RPE of 3-4 images minus the fluorescent signal in 3-4 no primary antibody control images, per mouse. Each data point represents one mouse. Mean +/- 95% C.I. are shown. Data were analyzed by 1-way ANOVA with Tukey’s multiple comparisons test. **** indicates p<=0.0001

This also suggests that the alternative pathway C3 convertase is primarily responsible for C3 activation and deposition at the RPE. Anti-C3d immunofluorescence appeared unaffected by *Abca4* genotype. Interestingly, anti-C3d labeling was still present in the choroid of *Cfd^-/-^* mice. This suggests that C3 convertase generated by the classical and/or lectin pathways is involved in C3 cleavage in the choroid but not RPE. In summary, while we did not observe any significant changes in C3 immunofluorescence with either antibody in *Abca4^-/-^* mice, loss of *Cfd* had a profound impact on C3d levels regardless of *Abca4* genotype indicating that alternative pathway is constitutively active in the RPE where it functions as the major driver of complement activity.

One caveat associated with the C3 immunofluorescence quantification is that C3 antibodies may collectively recognize different C3 cleavage fragments, some of which may be differentially regulated depending on genotype. We therefore performed western blotting using lysate prepared from 6–12-month-old mouse eyecup tissue (RPE, choroid, sclera) prepared under reducing conditions (Figure 4A). Under these conditions, the anti-C3 antibodies may recognize many different C3 α-chain fragments (Figure 4A), enabling quantification of specific C3 breakdown products based on known molecular weights. As an important caveat, the C3d region of the C3 protein contains the thioester site (35), which participates in covalent binding to target molecules and affects the molecular weight of opsonized C3b (36). Consequently, opsonized C3b, iC3b, C3dg and C3d α-chain fragments may not be detected at predicted molecular weights by western blot.

**Figure 4.**
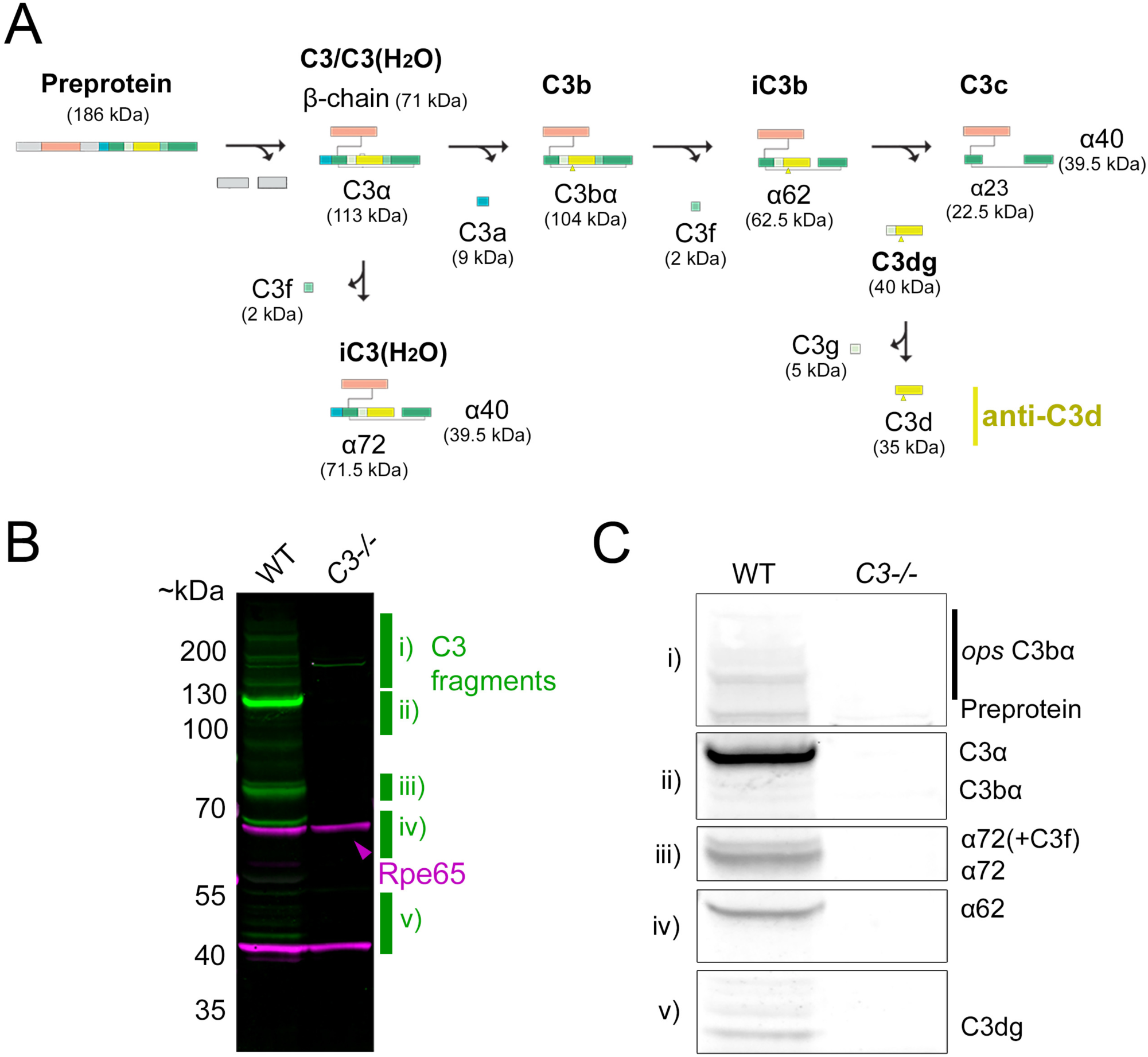
Qualitative identification of C3 fragments from wild type and *C3^-/-^* mouse eyecup tissue by western blot. A) Schematic representation of production and breakdown of the C3 protein. Estimated fragment molecular weights under reducing conditions are shown. Western blot was performed using eyecup lysate (RPE, choroid, sclera) detected using an antibody made against the C3d region (anti-C3d). Yellow shows the C3d region of the protein, which is part of the C3 α-chain (C3α). Note that the C3d region of the C3 protein contains the thioester site, which participates in covalent binding to target molecules (yellow arrowheads), potentially affecting fragment molecular weight of the C3b, iC3b, C3dg and C3d α-chain. B) Qualitative western blot images showing labelling of wild type and *C3^-/-^* eyecup tissue using the anti-C3d antibody. Marked regions i-vi indicate C3 fragments of interest and are shown zoomed-in in greyscale in panel (C) with predicted C3 fragment identities indicated.

First, we performed western blotting using the anti-C3d antibody to compare the pattern of C3 labelling between wild type and *C3^-/-^* eyecup lysate, identifying several known C3 protein fragments (Figure 4B,C). This antibody identified the C3 preprotein, C3/C3(H2O) α-chain fragment C3α, iC3(H2O) α-chain fragment α72 (37–39), iC3b α-chain fragment α62 and the C3dg α-chain fragment. The non-opsonized C3b α-chain fragment was not detected from wild type eyecup lysate using this antibody. Notably, none of these fragments were detected from the *C3^-/-^* eyecup lysate.

Next, we performed western blotting using the anti-C3d antibody and eyecup lysate from each experimental genotype, wild type, *Abca4^-/-^*, *Cfd^-/-^* and *Abca4^-/-^*; *Cfd^-/-^*(Figure 5A, full set of western blots used for quantification are in Supplementary Figure 5). In addition to the C3 fragments identified in Figure 4B,C, the anti-C3d antibody detected several large molecular weight fragments that were likely opsonized C3b, as described previously (36). The anti-C3d antibody also detected a suspected opsonized iC3b fragment at approximately 85 kDa, that was present at low quantities in *Cfd^+/+^* but absent from *Cfd^-/-^* eyecup lysate, mirroring our immunofluorescence results (Figure 3C).

**Figure 5.**
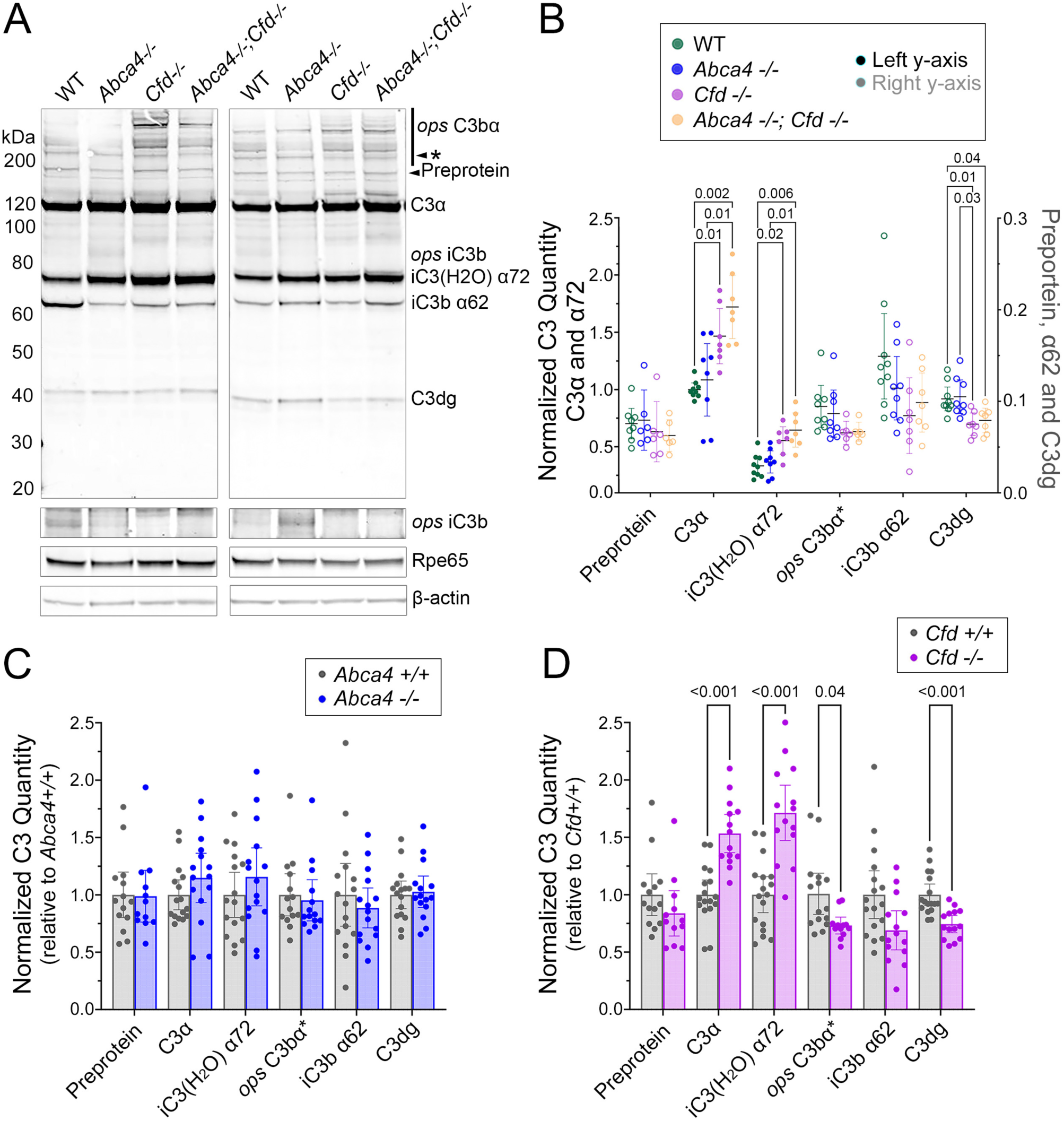
Quantification of C3 fragments from *Abca4^-/-^*, *Cfd^-/-^* mouse eyecup tissue by western blot. A) Representative blots labelled with the anti-C3d antibody, showing all genotypes. Note the faint C3 α-chain fragment detected with the anti-C3d antibody, which may be the opsonized iC3b α62 fragment (ops iC3b) and was frequently detected in *Cfd^+/+^*but not *Cfd^-/-^* conditions. Arrowhead (*) indicates the suspected opsonized C3b α-chain fragment selected for quantification. Higher molecular weight opsonized C3bα fragments were not quantified due to transfer inconsistencies and blot damage, which frequently affected the high molecular weight region of the blots. Rpe65 and β-actin were included as loading controls. B) Quantification of various C3 fragments under reducing conditions from mouse eyecup lysate, detected by the anti-C3d antibody. Animals used in this experiment range from 6-12 months of age. Relative quantities of C3 preprotein, C3α, iC3(H2O) α72, opsonized C3bα (*), iC3b α62 and C3dg were normalized to Rpe65 and β-actin, which were shown to have stable signal across the experimental conditions of interest (Supplementary Figure 6). Given the large number of experimental conditions, samples were run across multiple blots with a mix of genotypes per blot. To control for variability between blots, normalized C3 fragment quantities were expressed relative to the C3α fragment from WT samples. Note that quantities of C3α and iC3(H2O) α72 (filled circles) are represented on the left y-axis, and quantities of the preprotein, opsonized C3bα, iC3b α62 and C3dg fragments (empty circles) are represented on the right y-axis. Data were analyzed using a two-way ANOVA, revealing an effect of C3 fragment (p<0.0001), genotype (p<0.0001) and C3 fragment X genotype (p<0.0001). Multiple comparisons were performed using Tukey’s test, and p-values <0.05 are shown on the graph. C-D) Analysis of data from D, organized by independent variable (Abca4 genotype and Cfd genotype, respectively). Within each C3 fragment, data are represented relative to the respective control conditions (*Abca4^+/+^* or *Cfd^+/+^*). Data were analyzed using a two-way ANOVA, revealing an effect of C3 fragment X genotype for the *Cfd* genotype (p<0.0001) but not Abca4 genotype (p=0.6). Multiple comparisons were performed using Šídák’s test, and p-values <0.05 are shown on the graph. Each data point is a measurement from a single animal, and data are represented as mean +/- 95% C.I. for all graphs.

Finally, we performed quantitative western blotting using the anti-C3d antibody to measure the relative quantities of identified C3 fragments from wild type, *Abca4^-/-^*, *Cfd^-/-^* and *Abca4^-/-^*; *Cfd^-/-^*eyecup lysate (Figure 5B-D). C3 fragment quantities were expressed relative to both the RPE-specific protein Rpe65 and β-actin, which were present in stable quantities across the different conditions analyzed in this experiment (Supplementary Figure 6). We quantified bands corresponding to the C3 preprotein, C3/C3(H2O) α-chain, iC3(H2O) α72, presumptive opsonized C3b (ops C3bα*), iC3b α62 and C3dg. While many presumptive opsonized C3b large molecular weight fragments were detected (Figure 5A), we chose to quantify a single band running at approximately 200 kDa, which was reliably quantifiable across most samples. *Abca4* genotype had no clear effect on any C3 fragments (including comparison between wild type and *Abca4^-/-^*) quantified by western blot. In contrast to *Abca4*, quantities of the C3/C3(H2O) α-chain and iC3(H2O) α72 fragments appeared elevated in *Cfd^-/-^* relative to *Cfd^+/+^*, with a corresponding decrease in the relative quantities of activated C3 products, notably opsonized C3b and C3dg α- chain fragments. Additionally, neither genotype had a clear effect on the relative quantity of the C3 preprotein. Taken together, although we did not observe any changes associated with loss of *Abca4,* these data suggest that loss of *Cfd* reduces C3 convertase activity in the eyecup, resulting in decreased quantities of cleaved C3 fragments, such as opsonized C3b and C3dg, with a corresponding accumulation of non-activated C3 and iC3(H2O) fragments, C3α and α72.

Our final objective was to address alternative complement pathway involvement in the Stargardt mouse model ONL degeneration phenotype. However, *Abca4^-/-^*mice show minimal degeneration by 12-mo in albino strains (5,15,40) and no clear degeneration in pigmented strains (40,41). We initially planned to use *Abca4^-/-^;Rdh8^-/-^* double mutant mice as a model for rapid ONL degeneration as it has been reported to exhibit robust ONL degeneration by 3 months (28). In contrast, our testing (Supplementary Figure 7) and other studies (40) show no evidence of ONL degeneration in *Abca4^-/-^;Rdh8^-/-^*mice, suggesting that the initially characterized *Abca4^-/-^; Rdh8^-/-^* mice may have carried additional abnormalities that exacerbated ONL degeneration.

Given the absence of ONL degeneration in *Abca4^-/-^;Rdh8^-/-^*mice, we chose instead to use a bright light stress with the goal of accelerating ONL degeneration in our *Abca4^-/-^* mice. We harvested eye tissue from 6-12-mo wild type, *Abca4^-/-^*, *Cfd^-/-^* and *Abca4^-/-^;Cfd^-/-^* mice at 3- or 6 days post-light stress, and measured average ONL thickness in 250 µm increments along the length of the inferior and superior eyecup (Figure 6A,B). Across all genotypes, we observed consistent ONL thickness without and 3 days-post light stress, with an average central ONL thickness of 84 ± 8 µm without light stress thinning slightly to 65 ± 11 µm 3 days-post light stress (Figure 6C,D). In agreement with published data, we observed robust thinning of the ONL at 6 days-post light stress in the central retina of wild type mice that was partially rescued by loss of *Cfd*. Surprisingly, the loss of *Abca4* also appeared to be protective against light-induced ONL damage at 6 days-post light stress, similar to *Cfd^-/-^* mice, though the ONL thickness phenotype at 6 days-post light stress was highly variable between individuals within *Abca4^-/-^*, *Cfd^-/-^* and *Abca4^-/-^;Cfd^-/-^*mice. Light stress did not appear to affect ONL thickness in the peripheral retina and had no clear effect on C3 fragment quantities by WB (Supplementary Figure 8).

**Figure 6.**
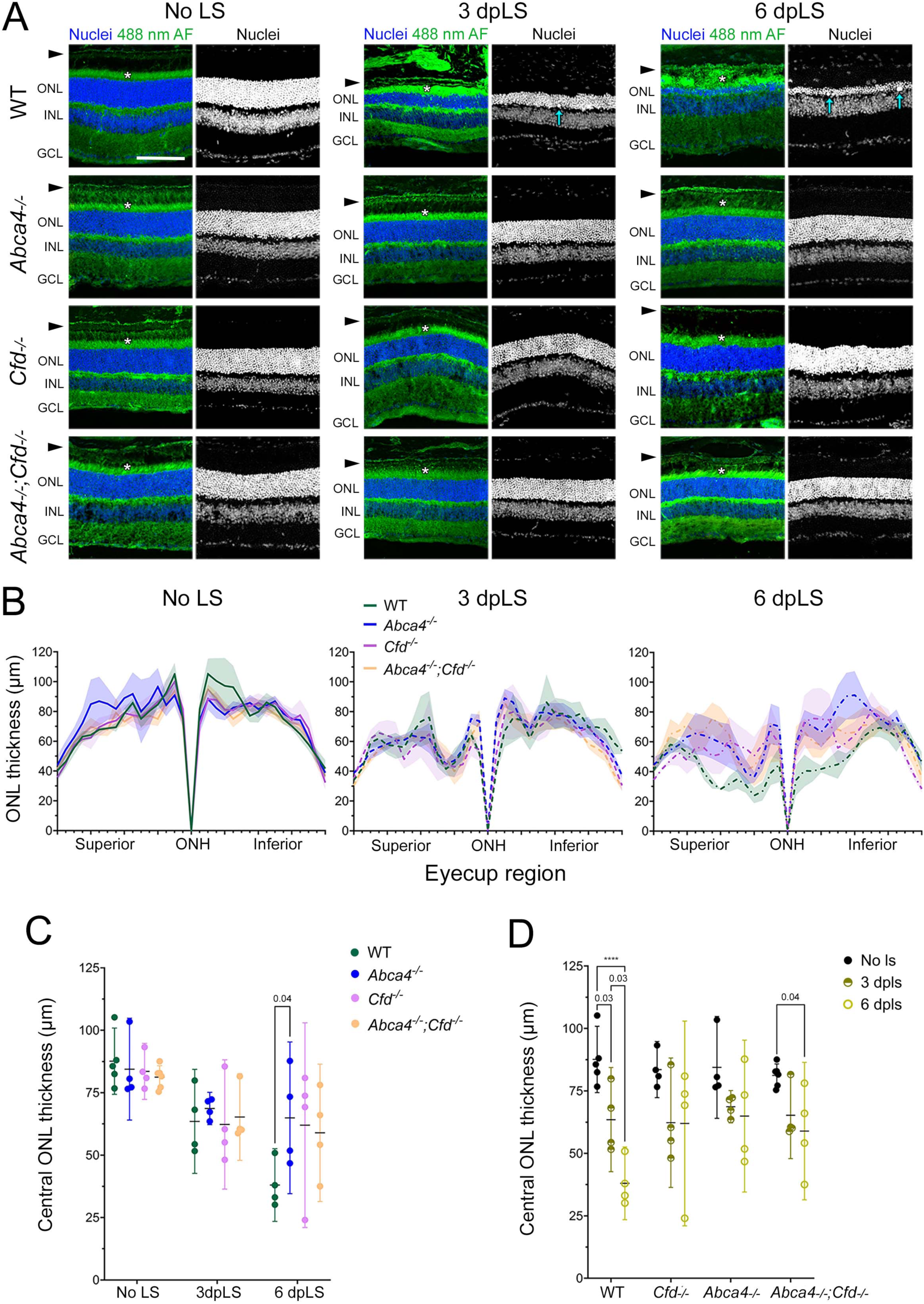
*Abca4^-/-^*and *Cfd^-/-^* mice show partial rescue of bright light-induced ONL degeneration. A) wild type, *Abca4^-/-^*, *Cfd^-/-^* and *Abca4^-/-^*;*Cfd^-/-^*mice were exposed to 20 klux bright for four hours to induce ONL degeneration. Eye tissue was harvested following either no light stress (LS) or 3- and 6-days post-LS (dpLS). Eye tissue was PFA-fixed and prepared as frozen sections in the inferior-superior orientation. Representative images show the superior region of the retina in the central eyecup adjacent to the optic nerve head (ONH). Blue shows DAPI nuclear labelling. Green shows 488 nm AF, enabling visualization of structures such as the inner and outer segments of the photoreceptors (*). Arrowheads indicate the RPE position and arrows indicate large swollen DAPI positive structures observed in the wild type ONL at 3 and 6 dpLS. Scale bar represents 100 µm. ONL, outer nuclear layer; INL, inner nuclear layer; GCL, ganglion cell layer. B) Average ONL thickness measured in 250 µm increments in the superior and inferior eyecup, starting at the ONH. Mean +/- SEM bands are shown. C-D) Average ONL thickness in the central half of the eyecup organized by genotype and LS condition, respectively. Each data point represents the thickness measured from a single mouse, averaged across two replicate tissue sections. Data represent mean +/- 95% CI. Data were analyzed by two-way ANOVA with Tukey’s multiple comparison’s test. **** indicates p<0.0001.

## Discussion

Complement system activation has been implicated as a contributing factor to the progression of the STGD1 phenotype, making its disruption a potential therapeutic strategy to slow or halt disease progression. To directly test the requirement of alternative pathway activation in a mouse STGD1 model, we employed a genetic rescue approach by generating mice lacking both *Abca4* and *Cfd*, the factor B–cleaving cofactor required for alternative pathway activation. Loss of *Cfd* markedly reduced C3d immunodetection in the RPE, both in the presence and absence of *Abca4*, but did not alter the elevated 488 nm autofluorescence characteristic of *Abca4^-/-^*mice. In a light-stress paradigm of photoreceptor degeneration, the absence of *Cfd* conferred a mild protective effect, and unexpectedly, a similar protection was observed in *Abca4^-/-^* mice. To our knowledge, this represents the first use of a direct genetic approach to investigate the role of complement activation in a STGD1 model.

Our findings contrast with those of Lenis et al. (2017), who reported that AAV-mediated expression of the complement inhibitor CRRY in the RPE of *Abca4^-/-^* mice robustly reduced 488 nm RPE autofluorescence and bisretinoid accumulation. It has previously been proposed that lipofuscin impedes clearance of C5b-9 via the endocytic pathway and lysosomal degradation, promoting a vicious cycle of complement attack that accelerates lipofuscin accumulation (15,42).

Although our data do not support this hypothesis, it is important to note that Lenis and colleagues used albino *Abca4^-/-^* mice, which undergo slow photoreceptor degeneration (40,43,44), whereas our experiments were performed in pigmented mice that do not exhibit photoreceptor loss (17,41,45). It is possible that pigmentation may act as an oxidative stress scavenger (40), mitigating A2E-epoxide–mediated damage and reducing complement activation to levels that do not drive lipofuscin accumulation. Consistent with this interpretation, we detected no changes in C3 breakdown products by immunolabeling or western blot in *Abca4*^-/-^ mice. This difference in background pigmentation may also explain our inability to replicate reports of altered C3 processing in *Abca4^-/-^* eyecups or RPE/choroid preparations (13,14,16,33). To further clarify the impact of pigmentation on complement activation and lipofuscin accumulation, we are currently generating *Abca4^-/-^;Cfd^-/-^* mice on an albino background.

Regardless of *Abca4* status, our findings indicate that CFD-mediated activation of the alternative pathway is the primary mechanism driving the generation of C3 breakdown products in the RPE. This is consistent with Williams et al (2016), who reported increased levels of full-length C3 and decreased levels of C3b/iC3b/C3c immunolabeling in the RPE of *Cfd^-/-^*mice.

Similarly, we observed elevated full-length C3 in eye cups by western blot and an almost complete loss of C3d immunolabeling in the RPE of *Cfd^-/-^* mice. Although the C3d antibody can detect intact C3 by western blot, our data suggest that in fixed tissue it does not recognize full-length C3 and instead preferentially detects downstream cleavage products that expose the His1002–Arg1303 epitope.

In wild-type RPE, abundant *Cfd*-dependent C3d immunolabeling localizes primarily to the basal labyrinth, with lower levels observed in apical microvilli and along cell boundaries. These findings support the idea that the alternative pathway actively surveys the RPE and contributes to cellular homeostasis. Whether this represents a non-canonical role for alternative pathway activation in maintaining RPE homeostasis remains unclear. However, the absence of an overt lipofuscin accumulation phenotype in *Cfd^-/-^* mice relative to wild type suggests that the complement system does not play a major role in regulating bisretinoid/A2E.

Bright white light–induced retinal damage has been widely used to produce ONL degeneration in both wild-type and disease-model mice (19,24,27–31). Using this paradigm, ONL degeneration was observed in treated relative to untreated *Abca4^-/-^* mice (31), though no comparison was made to wild type. Additionally, previous studies reported that albino *Abca4^-/-^* mice are more susceptible than wild-type mice to blue light–induced damage, specifically (47), and *Cfd^-/-^* mice exhibit reduced photoreceptor cell death following exposure to continuous, moderate-intensity white light (29). We therefore hypothesized that *Abca4^-/-^* mice would show *Cfd*-dependent photoreceptor degeneration greater than wild-type mice, using a bright light model. Unexpectedly, however, *Abca4^-/^-* mice displayed a mild protective effect. Reactive oxygen species generated by bright light are believed to drive degeneration, and blue light in particular promotes A2E photooxidation (8,48). Notably, in *Rpe65^-/-^*mice, photoreceptor degeneration is inhibited following both bright white light (24) and blue light (47) stress.

Furthermore, dark rearing prevents photoreceptor degeneration in mice lacking rhodopsin but interestingly not in mice lacking transducin, suggesting a non-canonical rhodopsin-dependent, transducin-independent mechanism (49). In *Abca4^-/-^* mice, impaired 11-cis-retinal (and rhodopsin dependent) regeneration may similarly confer partial protection.

Differences between our findings and those of Wu et al. (47), who observed exacerbated degeneration using a blue light stress model, may reflect the distinct mechanisms engaged by each paradigm. We selected a bright white light model to accelerate the degenerative processes characteristic of STGD1 while more closely approximating physiological light exposure. White light simultaneously drives 11-cis-retinal photoisomerization, exacerbating the at-retinal recycling defect in *Abca4^-/-^* mice, and promotes reactive oxygen species formation from accumulated A2E. In contrast, the blue light model primarily induces A2E-mediated RPE apoptosis (48), which may limit its ability to model the combined photoreceptor and RPE pathology that develops in STGD1. Nonetheless, our findings also indicate that the bright white light paradigm represents an acute injury distinct from the chronic, progressive degeneration in individuals with Stargardt disease.

## Acknowledgements

This work was funded in part by the National Sciences and Engineering Research Council of Canada (RGPIN-2024-06782) and the Canadian Institute for Health Research (PG 137101) to RLC. BCR was supported in part by a MITACS Postdoctoral Fellowship in partnership with Oak Bay Biosciences. KEG was supported by a National Sciences and Engineering Research Council of Canada Undergraduate Summer Research Award and University of Victoria Science Emerging Researcher Award. KEG and CNG were supported by the University of Victoria Jamie Cassels Undergraduate Research Award OS was supported by a University of Victoria Summer Undergraduate Research Award. We thank Courtney Gauthier and Chris Nelson for generously providing use of equipment and reagents and guidance with western blotting. We thank Roxana Radu, David Kroeger and Carl Erickson for helpful discussion and suggestions.

