## Supplementary material for "Loss of Complement Factor D suppresses alternative pathway activation but fails to reduce lipofuscin accumulation in the retinal pigmented epithelium of *Abca4^-/-^*mice": Suppl Figs and Legends

### Supplementary Figures and Figure Legends

**Supplementary Figure 1. Light stress exposure apparatus.** A) The light exposure apparatus used for this experiment consisted of two larger plastic storage containers lined with foil, into which four 1 L holding containers were fit. Centers of the holding container lids were cut and replaced with screen material, to keep the mice contained during light exposure while allowing air flow. A fluorescent light box containing 6 X 2 ft 144 W T4 fluorescent bulbs with 6500 K colour temperature (Grow Lights Canada, FLP26) was positioned above the holding containers to achieve 19-20 klux illumination inside the containers. Fans were used to maintain temperatures between 22-23°C during the experiment. B) Illumination was tested using a light meter (Extech instruments, LT300) with detector positioned facing upward inside a closed holding container prior to each experiment.

**Supplementary Figure 2. 488 nm autofluorescence is elevated in *Abca4*$^{-/-}$ RPE at 12-mo but not 6-mo.** A) Representative images showing 488 nm autofluorescence (AF) (green) in the RPE of 6-mo and 12-mo wild type and *Abca4*$^{-/-}$, taken from dark-adapted PFA-fixed paraffin embedded whole eye tissue sections. The basal (top) and apical (bottom) RPE boundaries are defined by dotted lines. Blue shows DAPI labelled nuclei. All images were collected using the same imaging settings. B) 488 nm AF histogram data showing the percentage of pixels in the RPE with different relative pixel intensities. Note that most pixels had a relative value of approximately 0 and are not included on the y-axis. Wild type samples had very few pixels above a relative intensity of 40. C) Plot of data from B showing the percent of pixels having a relative intensity greater than 40. D) Quantification of 488 nm AF from the same samples as C, instead measuring mean fluorescence intensity in the RPE. Note that both the threshold and mean fluorescence approaches to quantification of 488 nm AF produce similar results. Data for C and D were analyzed by one-way ANOVA ($p=0.0056$ and 0.0023, respectively) with Tukey's multiple comparisons and are represented as mean +/- 95% C.I. Each data point represents the average of eight images from a single mouse, imaged across two tissue sections, with four images collected immediately superior to the optic nerve head and four inferior.

**Supplementary Figure 3. Localization of C3 immunofluorescence to different RPE subcellular structures.** Validation of RPE subcellular structures labelled with the anti-C3d antibody (R&D Systems, AF2655) by co-labelling with either A) phalloidin, B) RPE65 or C) ZO-1. A) Phalloidin (ThermoFisher, A34055) detects F-actin and labeling was performed using wild type PFA-fixed frozen mouse eyecup tissue sections, showing that RPE C3d immunofluorescence is associated with the RPE basal labyrinth (arrows) and apical microvilli (*, AMV) cytoskeleton. Scale bar represents 20 µm. B) RPE65 (ThermoFisher, PA5-110315) immunolabelling on wild type PFA-fixed paraffin embedded mouse eyecup tissue sections is restricted to the RPE soma and absent from the apical microvilli (34), while RPE C3d immunofluorescence is associated with the apical microvilli (*, AMV) and strongly labels the RPE basal region (arrows). Scale bar represents 15 µm. C) ZO-1 antibody (ThermoFisher,61-7300) labelling was performed using wild type and *Cfd*$^{-/-}$ PFA-fixed paraffin embedded eyecup tissue sections and detects tight junctions between RPE cells (Jin et al., 2002), showing that C3d immunofluorescence is associated with lateral RPE membranes (arrowheads). Note that in the *Cfd*$^{-/-}$ mouse tissue, C3d immunofluorescence is lost in the RPE but still present in choroidal capillaries (CC). Scale bar represents 10 µm. Images also indicate Bruch's membrane (BrM), which frequently appears as a region of low AF between the RPE basal labyrinth and choroid, and the photoreceptor outer segments (OS).

**Supplementary Figure 4. Additional Quantitative C3 immunofluorescence data.** A) Quantification of C3d (R&D Systems) immunofluorescence in RPE basal labyrinth, cell body and apical microvilli from 12-mo paraffin sections. B) Quantification of total C3 (MP Biomedicals) immunofluorescence signal in the RPE and choroid, respectively, from 6-12-mo frozen sections. Animals were grouped based on *Cfd* genotype, irrespective of *Abca4* genotype. Total immunofluorescence was quantified by measuring the mean signal in the region of interest from 3-4 images minus the fluorescent signal in the region of interest from 3-4 no primary antibody control images, per mouse. Data were expressed relative to the $Cfd^{+/+}$ condition for each region. Each data point represents one mouse. Mean +/- 95% C.I. are shown. Data were analyzed by two-way ANOVA.

**Supplementary Figure 5. Summary of all blots used for quantification of C3 fragments from mouse eyecup tissue.** A and B) Blots used for quantification, showing C3 fragment labelling and Rpe65/b-actin labelling from light stress (LS) and 6-mo experimental samples. Eyecup tissue (RPE, choroid, sclera) was harvested from wild type, $Abca4^{-/-}$, $Cfd^{-/-}$ and $Abca4^{-/-};Cfd^{-/-}$ mice between about 6 and 12 months of age. $C3^{-/-}$ eyecup tissue was also harvested as a negative control. Tissue was collected as part of two experiments: A) a bright LS experiment where 6–12-month-old mice were either maintained in the dark or exposed to 20 klux for 4 hours, with tissue harvested three days post-LS, and B) an experiment where tissue was collected specifically at 6-mo without LS. Lysate was prepared from one eyecup per mouse under reducing conditions, and the same volume of lysate was loaded per lane of a gradient gel. A mix of different genotypes and experimental conditions were loaded per gel, with each gel containing multiple wild type samples without LS to serve as inter-blot controls. Blots were labelled using an anti-mouse C3d primary antibody (R&D Systems, AF2655) for quantification of different C3 fragments. Rpe65 and b-actin were used as loading controls. For a summary of C3 fragment sizes and fragments potentially recognized by this antibody, see the main figure.

Several known C3 fragments can be identified from these blots: the C3 preprotein at ~186 kDa, the C3 α-chain (C3α) at ~113 kDa, the iC3(H2O) fragment at ~72 kDa (α72), the iC3b α-chain fragment at ~62 kDa (α62) and the C3dg α-chain at ~40 kDa. The un-opsonized C3b α-chain (C3bα) is expected to run at ~104 kDa, at a slightly lower MW relative to the C3α, and was not reliably detected. Opsonized C3b fragments (C3b*, magenta) are predicted to run at a larger MW, and have been described previously (Law and Levine, 1977). A faint band smear at ~85 kDa (teal), likely corresponding to an opsonized iC3b α62 fragment (iC3b*), was also identified as being consistently present in *Cfd*$^{+/+}$ and absent from *Cfd*$^{-/-}$ samples, similar to RPE immunofluorescence data collected using this antibody.

We chose to quantify the C3 preprotein, C3α, α72, α62 and C3dg bands. Red indicates fragments and samples that were excluded from analysis due to low signal, incomplete transfer or blot damage.

**Supplementary Figure 6. Candidate loading control stability between experimental conditions of interest.** Given concerns about the possibility of RPE damage resulting from light stress, we first chose to assess the stability of the candidate loading controls in eyecup tissue harvested from wild type mice with and without exposure to retina-damaging LS. A) Blot images, showing Rpe65, β-actin and C3 α-chain bands used for quantification. Note placement of the MW ladder between wild type No LS samples 1 and 2. B) Inter-condition variability was measured by normalizing the signal for each LS sample to the mean No LS signal for each loading control.

We next reevaluated the stability of each candidate loading control across all planned experimental conditions. C) One sample per genotype (wild type, *Cfd$^{-/-}$*, *Abca4$^{-/-}$*, and *Abca4$^{-/-}$;Cfd$^{-/-}$*) without and with exposure to LS was included per blot, for a total of eight samples per blot. Four blots were performed in total, with one representative blot shown. D-F) Quantification of inter-condition variability for LS, *Abca4* genotype and *Cfd* genotype conditions, respectively.

Signal for Rpe65, β-actin and C3 α-chain bands was first normalized to a common inter-blot control sample within each blot. Data were grouped by independent variable and expressed relative to the control condition for each (No LS, *Abca4$^{+/+}$*, and *Cfd$^{+/+}$*). Inter-condition variability for each candidate loading control was evaluated by two-way ANOVA with Šídák's multiple comparisons test. P-values equal to 1 indicate no difference in signal between control conditions and experimental conditions. Smaller p-values indicate greater signal variability between conditions. Note that the C3 α-chain appears differentially regulated between *Cfd$^{+/+}$* and *Cfd$^{-/-}$* conditions (*). n=4 individual animals were used per condition, and each individual is represented by a single data point. Graphs show mean +/- 95% C.I.

Conclusions: Rpe65 and β-actin appear to produce reasonably stable signal between experimental conditions of interest, suggesting that they could be used in combination for performing lane normalization. The C3 α-chain should not be used for lane normalization, since it appears differentially regulated between *Cfd* genotypes.

**Supplementary Figure 7. No evidence of ONL cell death in the *Abca4$^{-/-}$;Rdh8$^{-/-}$* mouse retina at 6-10-mo.** A) Representative images showing H&E-stained wild type and *Abca4$^{-/-}$;Rdh8$^{-/-}$* retina. Images were collected superior and inferior to the optic nerve head (ONH). * shows ONL. B) ONL thickness in pixels measured from n=3 wild type and *Abca4$^{-/-}$;Rdh8$^{-/-}$* (DKO) mouse retinal sections. 0 indicates the position of the ONH. C) ONL thickness measured by optical coherence tomography (OCT) for wild type and DKO at different ages. 8 eyes (n=4 mice) were quantified for wild type samples, 16 eyes (n=8 mice) were quantified for 5-7 mo DKO, and 10 eyes (n=5 mice) were quantified for 8-10 mo DKO. Data in B and C show mean +/- SEM error bands. D) Density of DAPI positive nuclei in the ONL immediately superior and inferior to the ONH, comparing 6-mo wild type and DKO. One 8-mo DKO sample was included. Data represent mean +/- SD.

**Supplementary Figure 8. No clear effect of damaging LS on C3 fragment quantity by western blot.** 6-12-mo mice were exposed to 20 klux bright for four hours to induce ONL degeneration. Eyecup tissue (RPE, choroid, sclera) was harvested following either no light stress (LS) or 3 days post-LS for western blot. A) Quantification of various C3 fragments under reducing conditions from mouse eyecup lysate, detected by the anti-C3d antibody. Relative quantities of C3 preprotein, C3α, α72, opsonized C3b (C3b*), α62 and C3dg were normalized to Rpe65 and β-actin. Given the large number of experimental conditions, samples were run across multiple blots with a mix of genotypes per blot. To control for variability between blots, normalized C3 fragment quantities were expressed relative to the C3α fragment from wild type No LS samples. B) Analysis of data from panel A, organized by LS condition (No LS and 3 days post-4 hrs LS). Within each C3 fragment, data are represented relative to the No LS condition.

Data were analyzed using a two-way ANOVA, revealing a possible effect of LS condition ($p=0.01$). Multiple comparisons were performed using Šídák's test, showing specifically that the C3/C3(H2O) α-chain fragment may be elevated 3 days post-LS relative to No LS. C) Replication of C3α quantification using $n=4$ wild type mice without LS and 3 days post-LS reveals no clear effect of LS on the C3/C3(H2O) α-chain fragment.

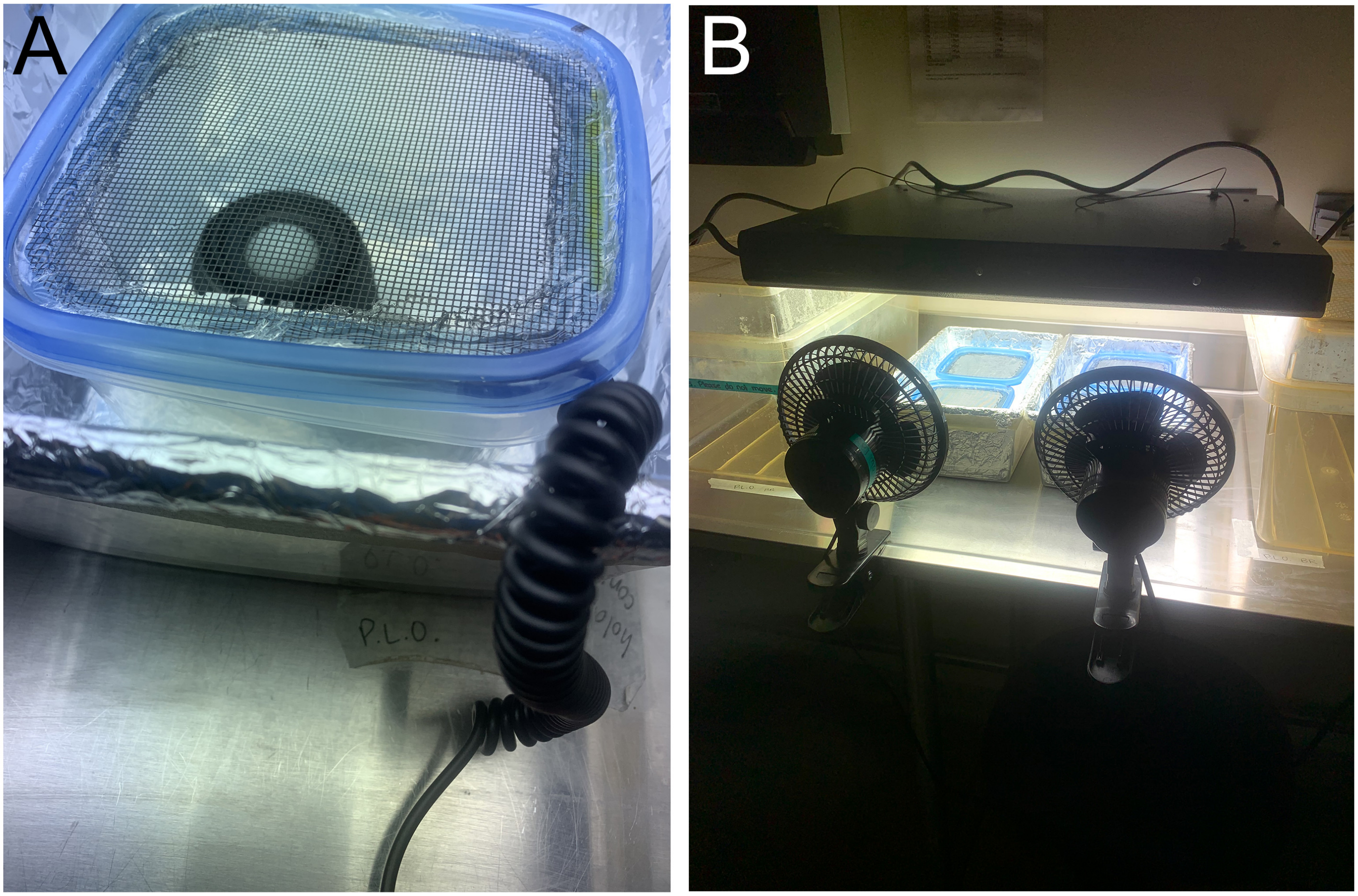


Suppl Fig 1

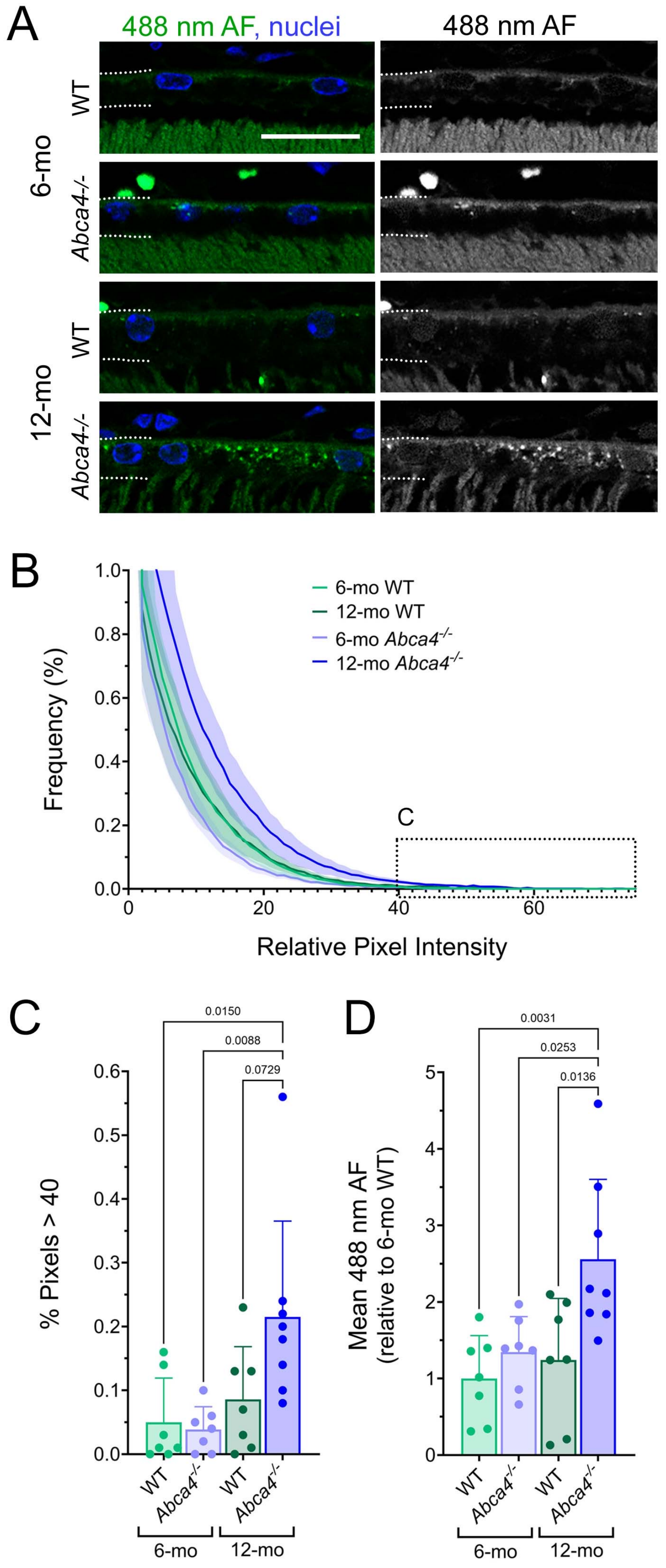
A
488 nm AF, nuclei
488 nm AF
6-mo
WT
Abca4-/-
12-mo
WT
Abca4-/-
B
Frequency (%)
Relative Pixel Intensity
6-mo WT
12-mo WT
6-mo Abca4-/-
12-mo Abca4-/-
C
C
% Pixels > 40
0.0150
0.0088
0.0729
WT
Abca4-/-
WT
Abca4-/-
6-mo
12-mo
D
Mean 488 nm AF (relative to 6-mo WT)
0.0031
0.0253
0.0136
WT
Abca4-/-
WT
Abca4-/-
6-mo
12-mo

Suppl Fig 2

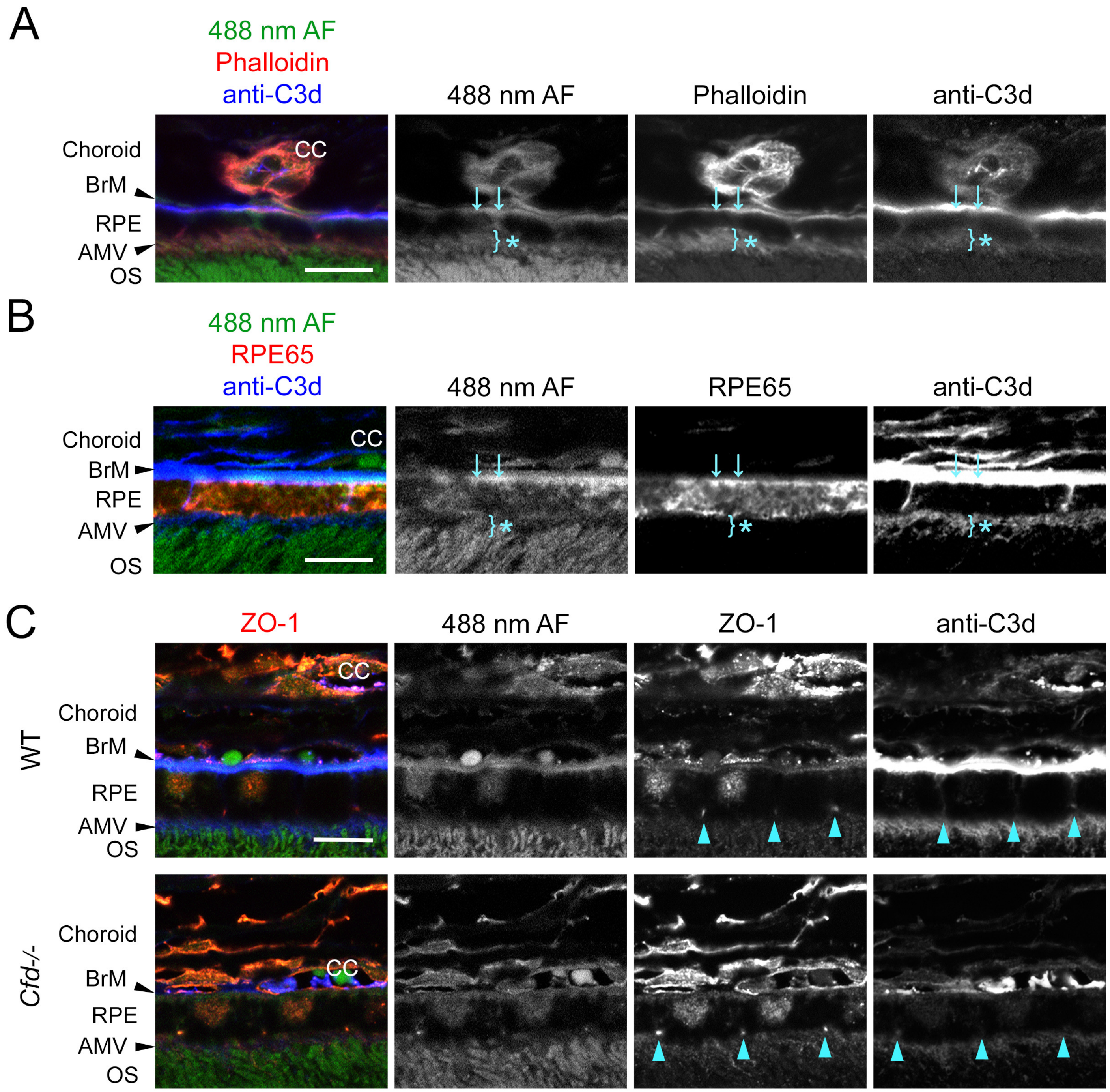

A
488 nm AF
Phalloidin
anti-C3d
488 nm AF
Phalloidin
anti-C3d
Choroid
BrM
RPE
AMV
OS
CC
B
488 nm AF
RPE65
anti-C3d
488 nm AF
RPE65
anti-C3d
Choroid
BrM
RPE
AMV
OS
CC
C
ZO-1
488 nm AF
ZO-1
anti-C3d
WT
Choroid
BrM
RPE
AMV
OS
CC
Cfd-/-
Choroid
BrM
RPE
AMV
OS
CC


Suppl Fig 3

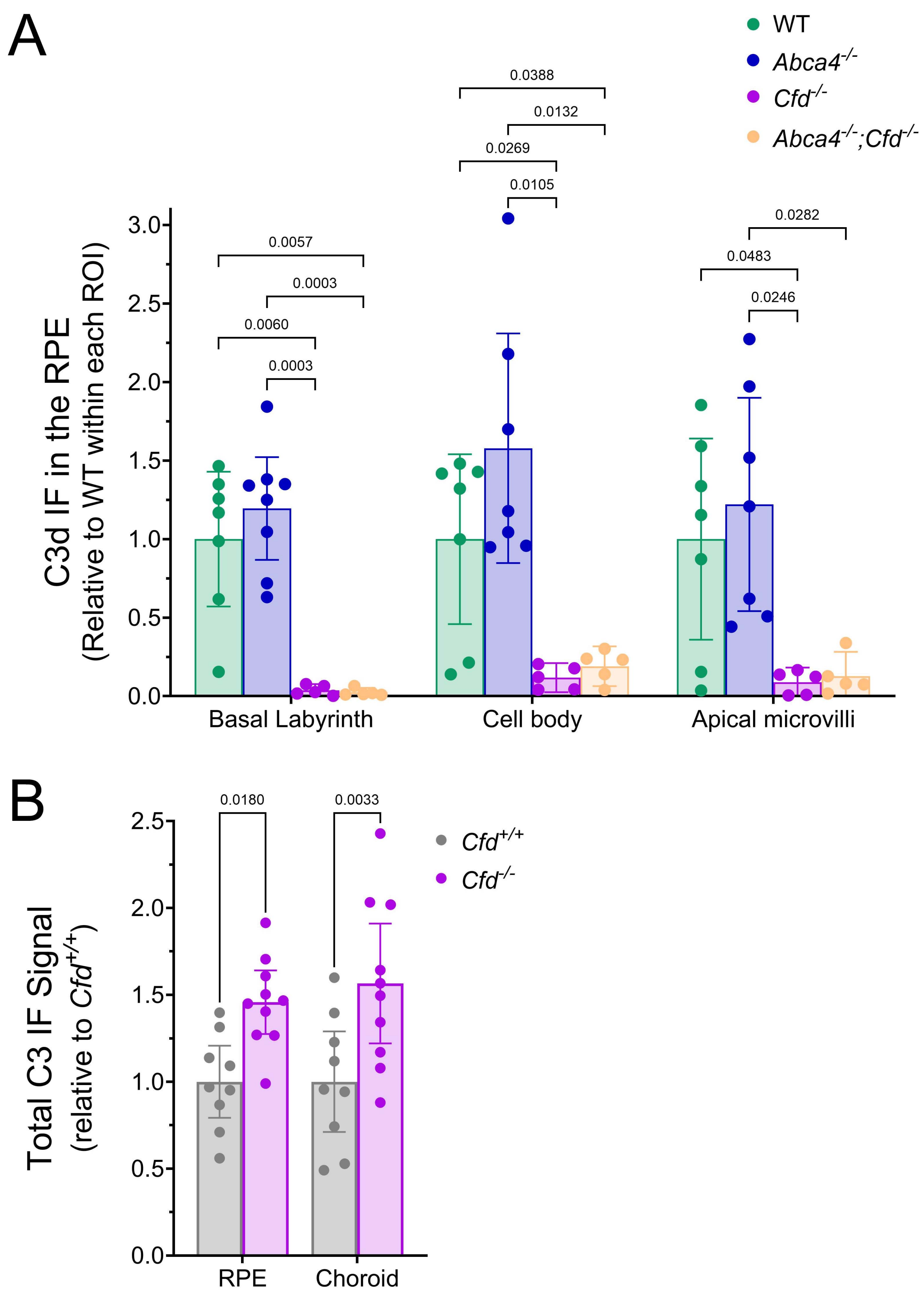
A
WT
Abca4-/-
Cfd-/-
Abca4-/-;Cfd-/-
C3d IF in the RPE
(Relative to WT within each ROI)
0.0388
0.0132
0.0269
0.0105
0.0057
0.0003
0.0060
0.0003
0.0282
0.0483
0.0246
3.0
2.5
2.0
1.5
1.0
0.5
0.0
Basal Labyrinth
Cell body
Apical microvilli
B
Total C3 IF Signal
(relative to Cfd+/+)
0.0180
0.0033
Cfd+/+
Cfd-/-
2.5
2.0
1.5
1.0
0.5
0.0
RPE
Choroid

Suppl Fig 4

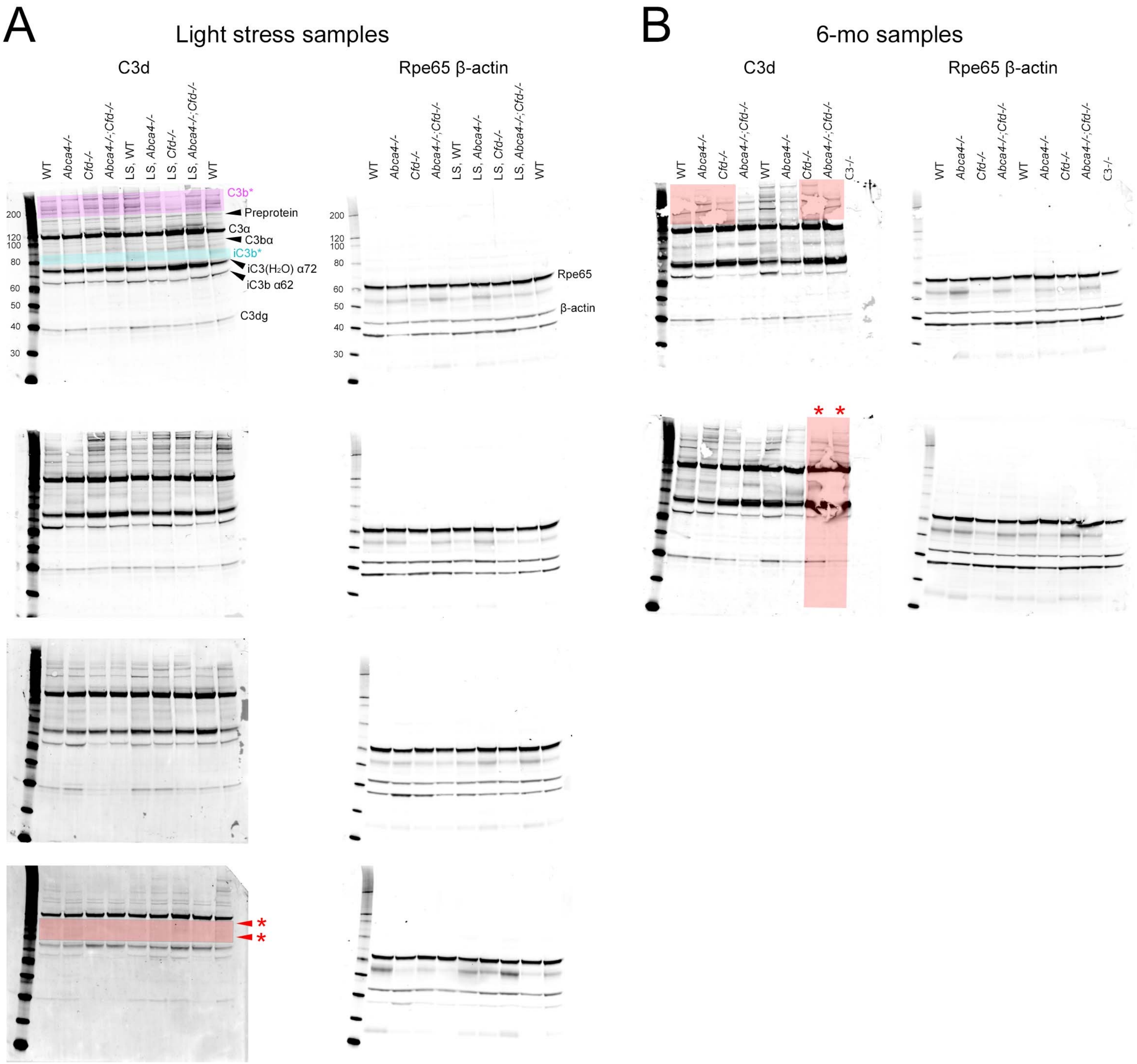

A
Light stress samples
C3d
Rpe65 β-actin
WT
Abca4-/-
Cfd-/-
Abca4-/-;Cfd-/-
LS, WT
LS, Abca4-/-
LS, Cfd-/-
LS, Abca4-/-;Cfd-/-
WT
C3b*
Preprotein
C3α
C3bα
iC3b*
iC3(H2O) α72
iC3b α62
C3dg
Rpe65
β-actin
200
120
100
80
60
50
40
30
Blot 1
Blot 2
Blot 3
Blot 4
B
6-mo samples
C3d
Rpe65 β-actin
C3-/-


Suppl Fig 5

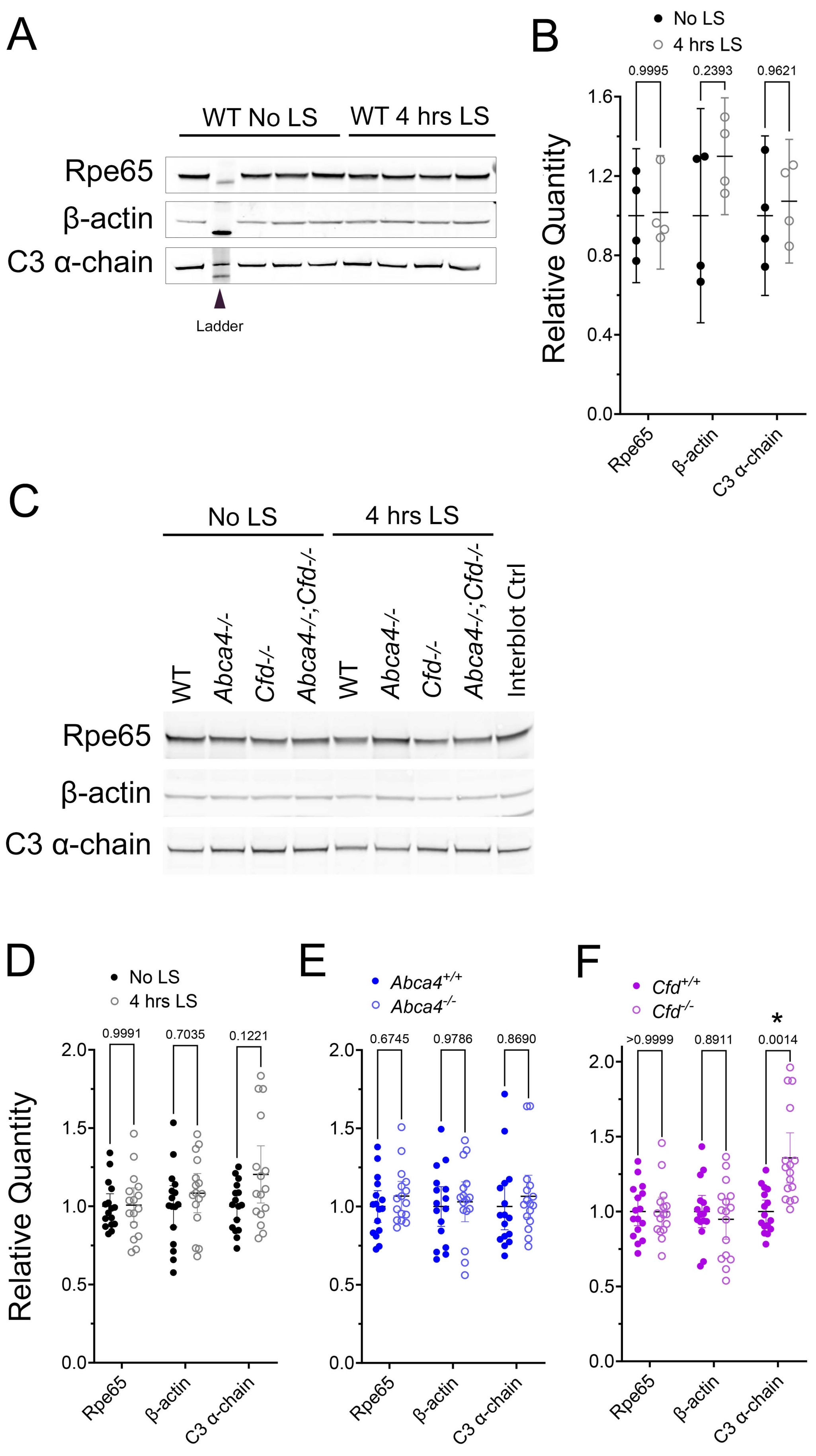
A
WT No LS
WT 4 hrs LS
Rpe65
β-actin
C3 α-chain
Ladder
B
No LS
4 hrs LS
0.9995
0.2393
0.9621
Relative Quantity
1.6
1.2
0.8
0.4
0.0
Rpe65
β-actin
C3 α-chain
C
No LS
4 hrs LS
WT
Abca4-/-
Cfd-/-
Abca4-/-;Cfd-/-
WT
Abca4-/-
Cfd-/-
Abca4-/-;Cfd-/-
Interblot Ctrl
Rpe65
β-actin
C3 α-chain
D
No LS
4 hrs LS
0.9991
0.7035
0.1221
Relative Quantity
2.0
1.5
1.0
0.5
0.0
Rpe65
β-actin
C3 α-chain
E
Abca4+/+
Abca4-/-
0.6745
0.9786
0.8690
2.0
1.5
1.0
0.5
0.0
Rpe65
β-actin
C3 α-chain
F
Cfd+/+
Cfd-/-
*
>0.9999
0.8911
0.0014
2.0
1.5
1.0
0.5
0.0
Rpe65
β-actin
C3 α-chain

Suppl Fig 6

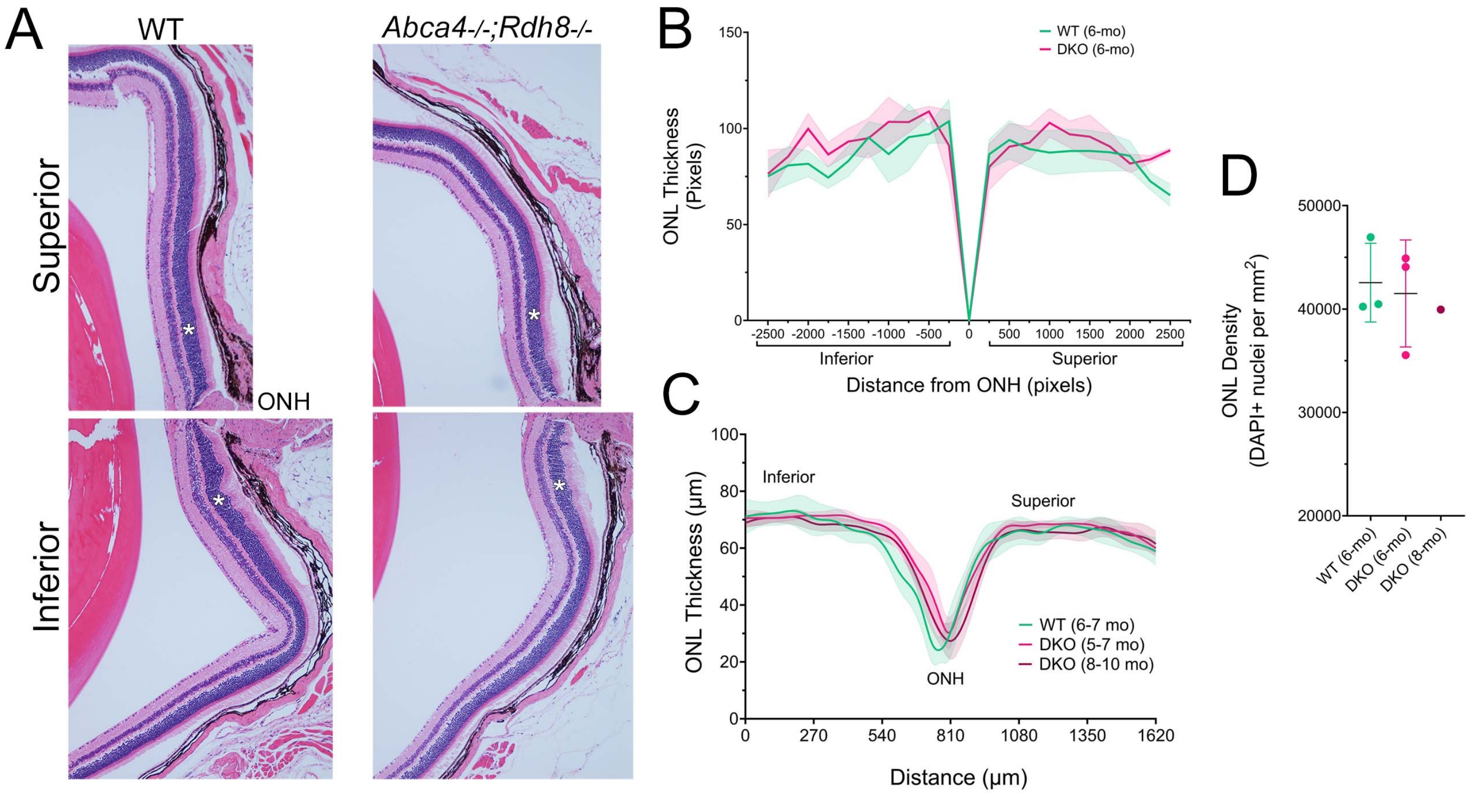
A
WT
Abca4-/-;Rdh8-/-
Superior
Inferior
ONH
B
WT (6-mo)
DKO (6-mo)
ONL Thickness (Pixels)
Distance from ONH (pixels)
Inferior
Superior
C
ONL Thickness (µm)
Distance (µm)
Inferior
Superior
ONH
WT (6-7 mo)
DKO (5-7 mo)
DKO (8-10 mo)
D
ONL Density (DAPI+ nuclei per mm2)
WT (6-mo)
DKO (6-mo)
DKO (8-mo)

Suppl Fig 7

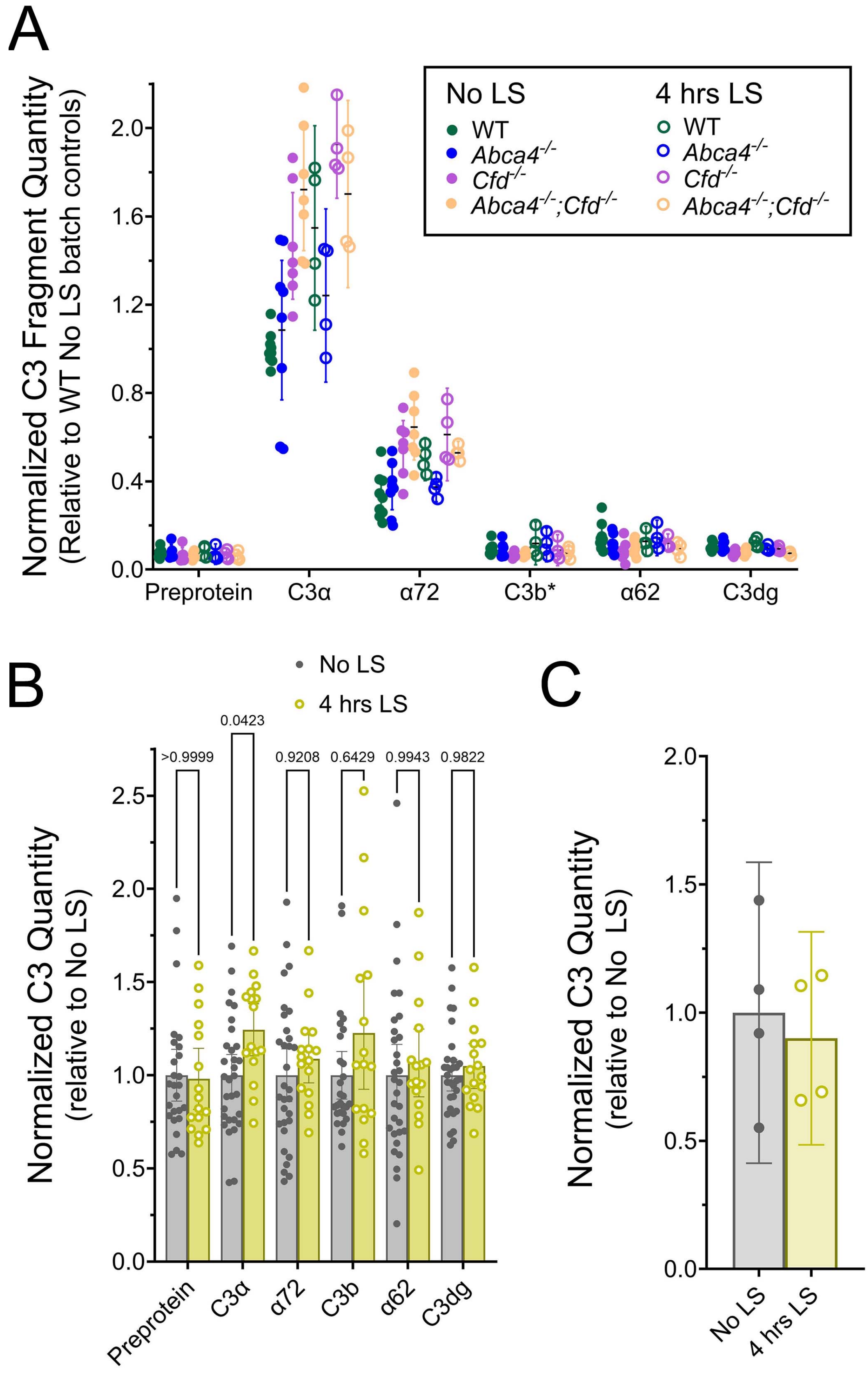
A
Normalized C3 Fragment Quantity
(Relative to WT No LS batch controls)
No LS
WT
Abca4-/-
Cfd-/-
Abca4-/-;Cfd-/-
4 hrs LS
WT
Abca4-/-
Cfd-/-
Abca4-/-;Cfd-/-
2.0
1.6
1.2
0.8
0.4
0.0
Preprotein
C3α
α72
C3b*
α62
C3dg
B
No LS
4 hrs LS
Normalized C3 Quantity
(relative to No LS)
>0.9999
0.0423
0.9208
0.6429
0.9943
0.9822
2.5
2.0
1.5
1.0
0.5
0.0
Preprotein
C3α
α72
C3b
α62
C3dg
C
Normalized C3 Quantity
(relative to No LS)
2.0
1.5
1.0
0.5
0.0
No LS
4 hrs LS

Suppl Fig 8
